# Tracing the evolution of single-cell cancer 3D genomes: an atlas for cancer gene discovery

**DOI:** 10.1101/2023.07.23.550157

**Authors:** Miao Liu, Shengyan Jin, Sherry S. Agabiti, Tyler B. Jensen, Tianqi Yang, Jonathan S. D. Radda, Christian F. Ruiz, Gabriel Baldissera, Moein Rajaei, Jeffrey P. Townsend, Mandar Deepak Muzumdar, Siyuan Wang

## Abstract

Although three-dimensional (3D) genome structures are altered in cancer cells, little is known about how these changes evolve and diversify during cancer progression. Leveraging genome-wide chromatin tracing to visualize 3D genome folding directly in tissues, we generated 3D genome cancer atlases of murine lung and pancreatic adenocarcinoma. Our data reveal stereotypical, non-monotonic, and stage-specific alterations in 3D genome folding heterogeneity, compaction, and compartmentalization as cancers progress from normal to preinvasive and ultimately to invasive tumors, discovering a potential structural bottleneck in early tumor progression. Remarkably, 3D genome architectures distinguish histologic cancer states in single cells, despite considerable cell-to-cell heterogeneity. Gene-level analyses of evolutionary changes in 3D genome compartmentalization not only showed compartment-associated genes are more homogeneously regulated, but also elucidated prognostic and dependency genes in lung adenocarcinoma and a previously unappreciated role for polycomb-group protein Rnf2 in 3D genome regulation. Our results demonstrate the utility of mapping the single-cell cancer 3D genome in tissues and illuminate its potential to identify new diagnostic, prognostic, and therapeutic biomarkers in cancer.

## Introduction

Cancer cells exhibit profound alterations in nuclear size, shape, and chromatin texture^1^. Microscopic examination of these structural features remains a gold standard for diagnosis and establishing pathologic cancer grade, which is frequently associated with prognosis^1^. It has become increasingly clear that the three-dimensional (3D) genome folding organization within the nucleus of cells plays a key role both in normal physiologic functions and in disease^2–9^. Here, we describe *in situ* genome-wide single-cell cancer 3D genome atlases, generated using an imaging-based 3D genomics method that we have pioneered – termed chromatin tracing^10,11^. Unlike earlier studies using sequencing-based 3D genomics technologies (e.g., high-throughput chromosome conformation capture (Hi-C)) that relied on population averaging of cells and indirect inference of altered genome organization in cancer cells^12–17^, our approach allows us to analyze 3D folding of the genome directly in individual cancer cells in the native tissue environment *in vivo*. In addition, it enables us to monitor evolution of this folding organization (and its regulation) during cancer progression from normal to preinvasive to invasive tumor cells. Our results define histologic stage-specific alterations in 3D genome architectures during lung and pancreatic adenocarcinoma (LUAD and PDAC) progression and illustrate how the evolution of 3D genome folding can be used to identify novel genes that govern prognosis, delineate cancer cell dependency, and regulate the 3D genome. Our findings and the methodologic toolkit we describe herein provide a unique resource by which *in situ* single-cell 3D genome information could be leveraged to identify potential new diagnostic, prognostic, and therapeutic cancer biomarkers.

## Results

### Genome-wide chromatin tracing in a mouse model of lung adenocarcinoma

To directly visualize chromatin folding organization in single cells within tissues, we performed genome-wide chromatin tracing in which we targeted a panel of 473 genomic loci spanning all 19 mouse autosomes at an average genomic interval of 5 Mb (**Supplementary Tables 1-3**). These included genomic regions that harbor “classic” oncogenes, tumor suppressor genes, and super-enhancers. To unambiguously distinguish genomic loci from one another, we adapted a previously published DNA multiplexed error-robust fluorescence *in situ* hybridization (DNA MERFISH) 100-choose-2 combinatorial barcoding design. We assigned each target genomic locus a unique 100-bit binary barcode, with each barcode containing two “1” bits and 98 “0” bits^18^; each “1” bit physically corresponded to a unique overhang readout sequence on the primary probes hybridized to the target genomic locus (**Fig. 1a**). The primary probes targeting each genomic locus contained two versions of readout sequences that corresponded to the two “1” bits in the barcode of the locus. We hybridized all primary probes to all target genomic loci and sequentially hybridized dye-labeled readout probes to image the 100 bits, with 50 hybridization rounds and two-color imaging (**Fig. 1a**). We then fitted the 3D positions of the foci imaged by fluorescence *in situ* hybridization (FISH), decoded the barcodes, and reconstructed chromatin traces in single cells (**Figs. 1b-1c**). To validate the quality of genome-wide chromatin tracing, we measured the mean spatial distance between each pair of target genomic loci and constructed a mean inter-loci distance matrix (**Fig. 1d**). We first performed this in lung alveolar type 2 (AT2) cells – the putative cell-of-origin for lung adenocarcinoma (LUAD)^19–21^ in wild-type (WT) mice. The resulting matrix featured lower *intra*-chromosomal distances and higher *inter*-chromosomal distances, consistent with chromosome territory organization^22^. Importantly, the inter-loci distances measured from WT AT2 biological replicates were highly correlated with each other (Pearson correlation coefficient = 0.91; **Extended Data Figs. 1a-1b**). Within each chromosome, the inter-loci spatial distances scaled with expected genomic distances, with longer chromosomes reaching larger spatial distances (**Extended Data Fig. 1c**). On average, inter-loci spatial distances followed a power-law function in comparison to genomic distances, with a scaling factor of about one-tenth (**Extended Data Fig. 1d**), similar to scaling factors calculated based on previous reports in other mouse tissues (**Extended Data Figs. 1e-1f**)^23,24^.

**Fig. 1.**
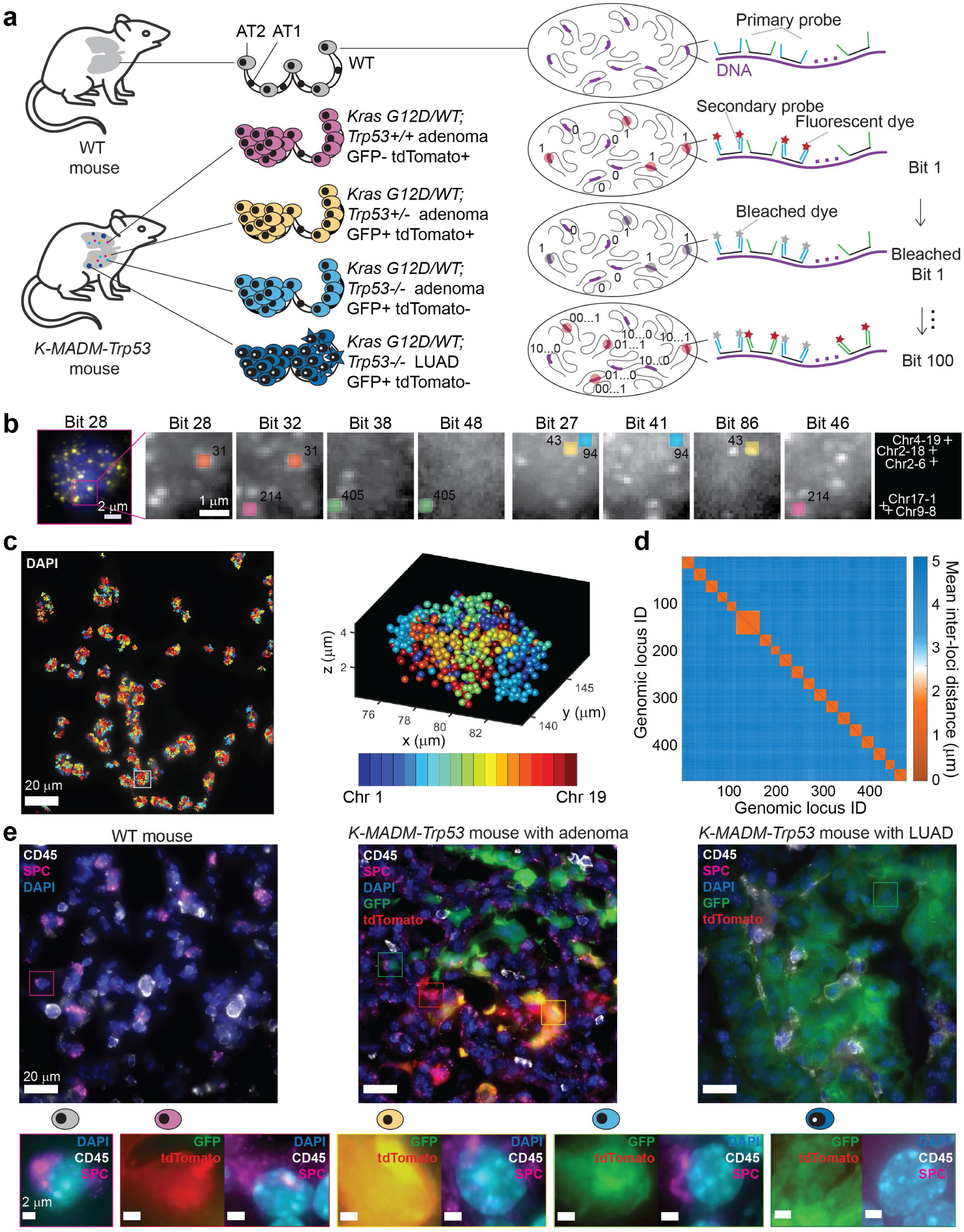
A genome-scale chromatin tracing strategy to visualize cancer 3D genomes. **a,** Schematic illustration of the experimental procedure. **b,** (Left panel) Raw FISH foci in bit 28 in a WT AT2 cell. (Right panels) Zoom-in images of raw FISH foci showing the decoding procedure and reconstructed genomic loci for 5 loci (shown as dots with 5 pseudo-colors each appearing twice). **c,** (Left panel) Reconstructed chromatin traces superimposed with DAPI staining. The traces are 2D projections of x, y coordinates. The DAPI image is a maximum-intensity z-projection from a 10-μm z-stack. (Right panel) The 3D positions of all decoded genomic loci in a single-cell nucleus. Different pseudo-colors represent different autosomes. **d,** Matrix of mean inter-loci distances between all genomic loci in AT2 cells. n = 4,806 AT2 cells. **e,** (Upper panels) Immunofluorescence staining of cell-type markers, DAPI staining, and fluorescent protein imaging of lung tissue from a WT mouse, a *K-MADM-Trp53* mouse with adenomas, and a *K-MADM-Trp53* mouse with LUAD. (Lower panels) Representative cells of each state. The images are maximum-intensity z-projections from 10-μm z-stacks.

Having established the approach in normal tissues, we next performed chromatin tracing on the *K-MADM-Trp53* mouse lung cancer model (**Fig. 1a, Extended Data Figs. 1g and 1h**) that induces sparse and sequential mutagenesis of *Kras* and *Trp53* in the lung epithelium to mimic the faithful genetic and histologic progression of human LUAD development^25–27^. To trace the evolution of 3D genome organization during LUAD progression in these mice, we leveraged the **<u>M</u>**osaic **<u>A</u>**nalysis with **<u>D</u>**ouble **<u>M</u>**arkers (MADM) system^28^, which allows unambiguous coupling of genotype to fluorescence labeling^28,29^. In this system, sibling green-fluorescent protein-expressing (GFP+) homozygous mutant and tdTomato-expressing (tdTomato+) homozygous WT cells are generated from a rare Cre/loxP-mediated *inter*-chromosomal mitotic recombination (G2-X) event in a heterozygous parent. Yellow heterozygous (GFP+/tdTomato+) cells are also generated through non-mitotic inter-chromosomal recombination (G0/G1) or mitotic recombination followed by Z segregation^28,29^. Subclones of green (GFP+) *Trp53*^−/−^, red (tdTomato+) *Trp53^+/+^*, and yellow (GFP+/tdTomato+) *Trp53^+/−^* cells can then be traced in preinvasive *Kras* mutant lung adenomas initiated by inhaled lentiviral Cre infection^30^. Only *Trp53*^−/−^, and not *Trp53^+/+^* or *Trp53^+/−^*adenoma cells, progressed to advanced LUAD^30^, providing us with a tractable *in vivo* model to study histologically defined cellular stages of lung cancer progression from normal AT2 to preinvasive adenoma to LUAD cells in lung tissue sections. We combined multiplexed fluorescence imaging of GFP, tdTomato, surfactant protein C (SPC), and CD45 with genome-wide chromatin tracing on the same tissue sections to achieve cancer-state-specific single-cell 3D genome profiling (**Extended Data Fig. 1g**). SPC marks AT2 and cancer cells, whereas the pan-immune marker CD45 allowed us to exclude macrophages, which are labeled alongside cancer cells by the MADM system (presumably through lentiviral Cre infection) (**Fig. 1e**). To build a complete 3D genome atlas that captures the full spectrum of tumor development, we analyzed 26,852 cells across 17 replicates collected from multiple lung lobes (most containing several discrete tumors) of 13 mice harboring all stages of LUAD progression: AT2, AdenomaR (red *Trp53^+/+^*), AdenomaY (yellow *Trp53^+/-^*), AdenomaG (green *Trp53*^−/−^), and LUAD cells.

### A structural bottleneck of the 3D genome in early lung cancer progression

Given the heterogeneous nature of cancer development^31^, we hypothesized that chromatin conformations might become increasingly diverse during subclonal progression. To our surprise, chromatin folding conformations instead became less heterogeneous during early progression from AT2 to preinvasive adenoma cells – quantified as the coefficient of variation (COV) of inter-loci distances along each autosome (**Figs. 2a, 2b, and Extended Data Fig. 2a)**. This trend was then reversed upon progression to LUAD. To define the dynamics of 3D genome alterations during LUAD progression, we quantified the population-averages for several structural features of chromosomes *in situ* for each cell state (AT2, Adenoma (R, Y, G), LUAD), including: (1) *intra*-chromosomal compaction (mean inter-loci distances on each autosome), (2) long-range intermixing (calculated using a demixing score metric benchmarked using published active and inactive X-chromosome folding conformations in human cells^11^) (**Extended Data Fig. 2b**), (3) active (A) and inactive (B) chromatin compartment polarization (quantified using a polarization index metric^11^), (4) *inter*-chromosomal distance, and (5) radial localization of each genomic locus in the nucleus (see **Methods**). *Increased* chromosome compaction was seen for all adenoma cell states compared with AT2 cells (decreased decompaction score; **Figs. 2a, 2c, and Extended Data Fig. 2c**), alongside *reduced* long-range intermixing (increased demixing score; **Fig. 2d and Extended Data Fig. 2d**) and *increased* A-B compartment polarization (**Fig. 2e**). Moreover, chromosomes in adenoma states were more spatially associated (reduced *inter*-chromosomal distance; **Fig. 2f and Extended Data Figs. 2e-2g**), with a subset (*e.g.,* Chr 7, 13, and 19) showing consistent changes in radial distribution within the nucleus compared to AT2 cells (**Fig. 2g and Extended Data Fig. 2i**). Strikingly, these phenotypes again largely reverted in LUAD cells (**Fig. 2 and Extended Data Fig. 2**). Importantly, *inter*-chromosomal distances in LUAD largely surpassed those of AT2 cells (**Fig. 2f and Extended Data Fig. 2h**), leading to an expected larger overall nuclear volume in LUAD than in AT2 and adenoma cells (**Extended Data Fig. 2j**). Together, these data demonstrate stereotypical, non-monotonic, and stage-specific 3D genome conformations during lung cancer progression, which could represent a structural bottleneck for tumor development.

**Fig. 2.**
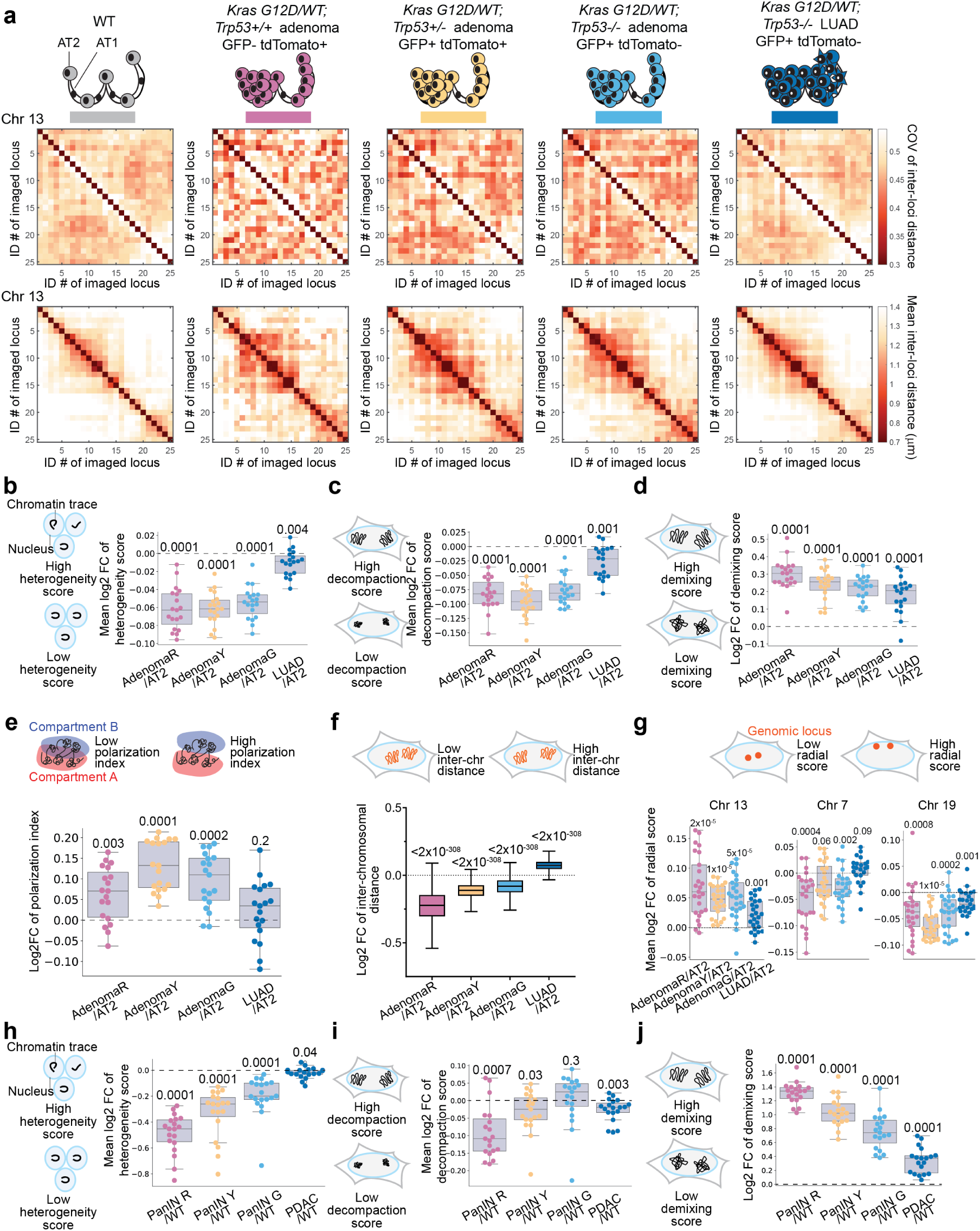
Systematic changes of 3D genome conformations during lung and pancreatic cancer progression. **a,** Coefficient of variation (COV) of inter-loci distances (upper panels) and mean inter-loci distances (lower panels) of mouse Chr13 in WT AT2 (n = 4,806), *Trp53^+/+^*adenoma (AdenomaR, n = 791), *Trp53^+/−^* adenoma (AdenomaY, n = 1,603), *Trp53^−/−^* adenoma (AdenomaG, n = 1,941), and *Trp53^−/−^*LUAD (n = 17,711) cells. Cell numbers of each cell state are identical in **a-g**. **b-e,** Distributions of the (mean) log2 fold change of heterogeneity (COV of inter-loci distance), decompaction (mean inter-loci distance), demixing scores, and polarization indices of each autosome (n = 19), comparing each cancer state to AT2 state. **f,** Distribution of the log2 fold change of inter-chromosomal distances, comparing each cancer state to AT2 state. **g,** Distribution of the mean log2 fold change of radial scores of Chr13, Chr7, and Chr19, comparing each cancer state to AT2 state. **h-j,** Distributions of the (mean) log2 fold change of heterogeneity (COV of inter-loci distance), decompaction (mean inter-loci distance), and demixing scores of each autosome (n = 19), comparing each pancreatic cancer state to normal duct cells. n = 1529, 123, 189, 361, 475 for normal duct, PanIN R, PanIN Y, PanIN G, and PDAC cells. Cell numbers of each cell state are identical in **h-j**. p values of two-sided Wilcoxon signed-rank test (**b-j**) are displayed. In **b-j**, the horizontal lines of each box from top to bottom represent the 75th percentile, median, and 25th percentile. Whiskers extend to the non-outlier maximum and non-outlier minimum. Outliers are defined as values at least 1.5 times interquartile range away from the top or bottom of the box.

We also identified several cancer state-independent 3D genome features. For example, spatial proximity frequencies were the highest for *trans*-chromosomal A-A compartment interactions, followed by A-B and B-B interactions (**Extended Data Figs. 3a-3i**). The same trend was also observed in long-range *cis*-chromosomal compartment interactions (**Extended Data Figs. 3j-3l**). Furthermore, we identified a consistent negative correlation between radial scores and A-B compartment scores (**Extended Data Fig. 3m**). These observations align with previously reported findings in mouse fetal livers and human lung fibroblast cells^18,24^, suggesting features that represent general principles of 3D genome organization independent of cell state, cell type, and even mammalian species.

### Structural features of 3D genome evolution are largely conserved in pancreatic adenocarcinoma progression

To study whether the global 3D genome changes observed in LUAD progression are conserved in other cancer types, we also leveraged a mouse model of pancreatic ductal adenocarcinoma (PDAC). We used an analogous *K-MADM-Trp53* model that directs Cre recombinase activity to the pancreas using a *Pdx1-Cre* transgene and recapitulates the major genetic and histologic properties of human PDAC progression^30,32,33^. We performed genome-wide chromatin tracing targeting the same 473 genomic loci combined with cytokeratin 19 (CK19; a marker for normal and malignant duct cells) immunofluorescence, GFP, tdTomato, and DAPI imaging in the same single cells. We analyzed a total of 2,677 cells at all stages of PDAC progression, including normal CK19+ duct cells, preinvasive pancreatic intraepithelial neoplasia (PanIN) cells (R, Y, G), and invasive PDAC cells, all in their native tissue microenvironment (**Extended Data Figs. 4a-4d**). Mean inter-loci distances showed the expected chromosome territory features and genomic distance scaling (**Extended Data Figs. 4e and 4f**), consistent with high-quality traces. Analysis of key architectural features of chromosomes *in situ* largely recapitulated findings in LUAD progression, including (1) *intra*-chromosomal heterogeneity (**Fig. 2h**), (2) *intra*-chromosomal compaction (**Fig. 2i**), (3) long-range intermixing (**Fig. 2j**), (4) *inter*-chromosomal distances (**Extended Data Fig. 4g**) and (5) radial localizations of genomic loci in the nucleus (**Extended Data Fig. 4h**). When comparing PanIN to WT duct cells, we observed that chromosomes largely exhibited decreased heterogeneity, more compact folding, reduced long-range intermixing, decreased *inter*-chromosomal distances, and changes in radial distribution for a subset of chromosomes (*e.g.,* Chr 1, 17, and 19). These phenotypes were at least partially reverted upon progression to PDAC, except for *inter*-chromosomal distances. Thus, stereotypical, non-monotonic, and stage-specific changes in 3D genome conformations seen in PDAC progression mirrored those seen in lung cancer, arguing that they may represent general features of tumorigenesis.

### Organization of the 3D genome reflects specific histologic cancer cell states at the single-cell level

We next asked whether it is possible to classify the different histologic cancer cell states in our models based solely on the single-cell 3D genome data – defining state-specific *in situ* single-cell biomarkers. We first established, using subsampling analysis, that as few as 100 cells per cell state are sufficient to recapitulate 3D genome organization changes for most structural features described above (**Extended Data Figs. 5a-5f**). We then leveraged a “Trace2State” pipeline that performs dimensionality reduction of single-cell 3D genome conformation data to visualize clusters corresponding to different histologic cell states in a low dimensional representation. We adopted a previously developed single-cell A/B compartment (scA/B) score metric for all target genomic loci (**Fig. 3a**)^34,35^. The scA/B score of each observed locus is defined as the mean A/B compartment score of all its spatially adjacent loci (<1200-nm radius) and reflects the A/B compartment identity of each locus at the single-cell level. We used these scores as high-dimensional input variables to generate a two-dimensional display using three independent dimensionality reduction methods^36–38^. As shown in **Fig. 3a**, the single-cell 3D genome conformations of AT2 (purple) and LUAD (blue) cells were largely distinct in these plots, whereas those of preinvasive adenoma cells were clustered between these populations, concordant with histologic progression. We trained a series of supervised machine learning models to classify the different cancer cell states based on the single-cell 3D genome data (see **Methods**). A support vector machine model performed best, with 90% overall accuracy in predicting cancer cell states (**Figs. 3b-3c**). Similar results were obtained for the PDAC data (**Figs. 3d-3f**). These results argue that analysis of 3D genome organization can be used to distinguish between histologic cell states during oncogenesis at the single-cell level.

**Fig. 3.**
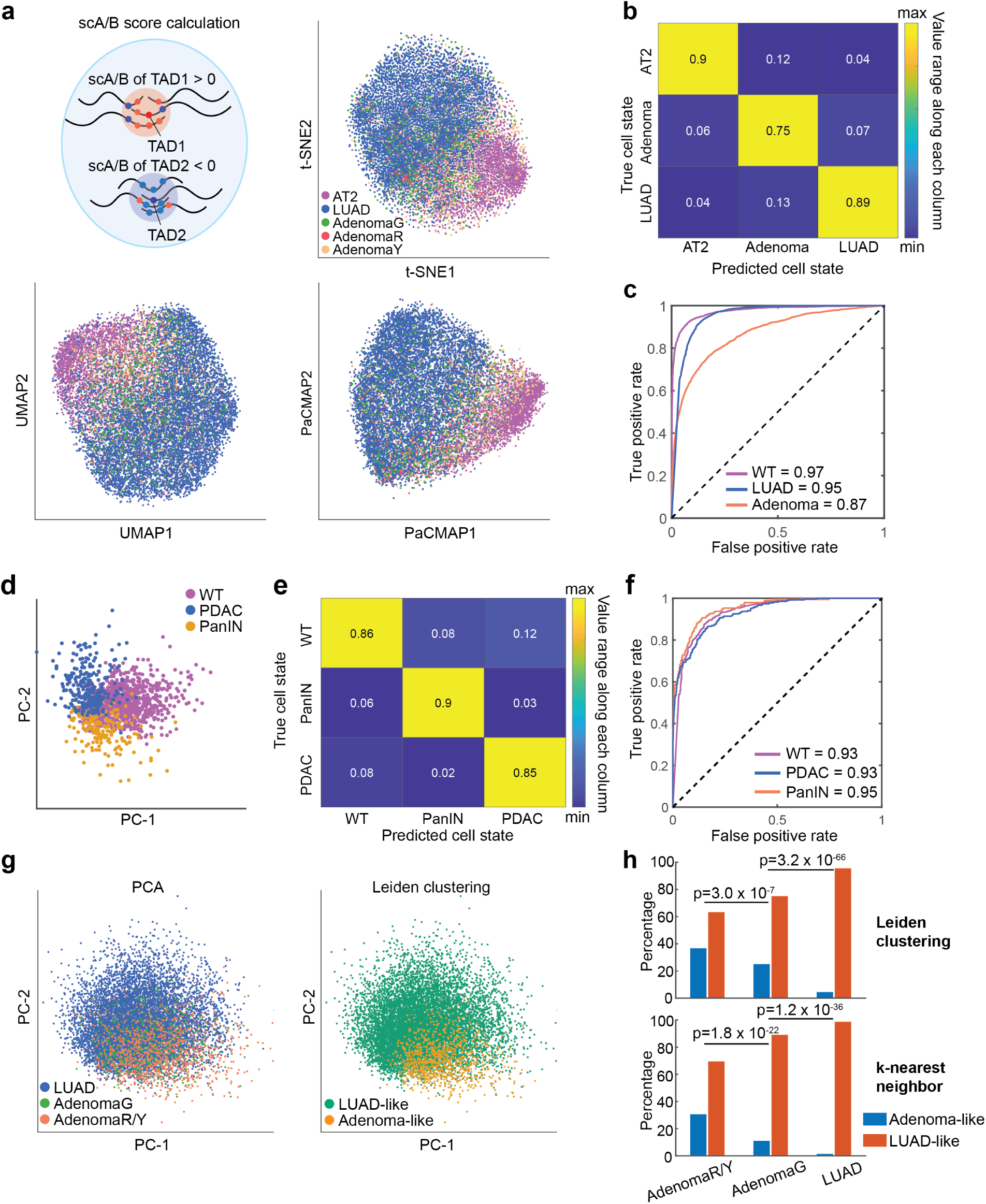
The single-cell 3D genome distinguishes and encodes cancer progression states. **a,** Cartoon illustration of scA/B score calculation (left panels). t-SNE, UMAP and PaCMAP plots of single-cell 3D genome conformations (right panels). n = 3410, 157, 689, 878, and 8834 for WT AT2, AdenomaR, AdenomaG, AdenomaY, and LUAD cells, respectively. Cell numbers of each cell state are identical in **a-c**, **g** and **h**. **b,** Confusion matrix of supervised machine learning in mouse lung cells. The number in each matrix element represents the precision in each predicted state. **c,** Receiver operating characteristic (ROC) curves of the machine learning model in mouse lung cells. The area under curve (AUC) values are shown. **d,** PCA plot of single-cell 3D genome conformations. n = 1103, 191, and 268 for normal duct, PanIN, and PDAC cells, respectively. Cell numbers of each cell state are identical in **d-f**. **e,** Confusion matrix of supervised machine learning in mouse pancreas cells. The number in each matrix element represents the precision in each predicted state. **f,** Receiver operating characteristic (ROC) curves of the machine learning model in mouse pancreas cells. The area under curve (AUC) values are shown. **g,** PCA plot of single-cell 3D genome conformations of adenoma and LUAD cells (left). Leiden clustering separates Adenoma-like and LUAD-like clusters (right). **h,** Percentages of cells with adenoma-like or LUAD-like 3D genome conformations in **g**, in each of the AdenomaR/Y, AdenomaG, and LUAD states. The adenoma-like or LUAD-like conformation state for each cell is assigned based on a Leiden clustering approach (upper) or the majority state of its five nearest neighbors (lower) in the left panel of **g**. p values from two-sided Fisher’s exact test are shown.

We hypothesized that the subset of *Trp53^−/−^* AdenomaG cells that progress into LUAD without clear mutational drivers (**Extended Data Figs. 6a-6c**) might have acquired specific 3D genome features associated with this behavior. Indeed, AdenomaG cells often exhibited chromosome-scale structural features intermediate between those of non-progressing AdenomaR/Y cells and LUAD cells (**Figs. 2b-2d**). 3D genome dimensionality reduction analyses further showed that preinvasive AdenomaR/Y cells and invasive LUAD cells formed distinct but overlapping clusters, with AdenomaG cells dispersed within and between them (**Fig. 3g**). Using either a Leiden clustering approach^39^ or a k-nearest-neighbor method, we computationally segmented the low-dimensional space into two regions of adenoma-like and LUAD-like conformations and quantified the percentage of AdenomaG cells distributed in the two regions. In this analysis, a greater proportion of AdenomaG cells (vs. other adenoma cells) adopted LUAD-like 3D chromatin organization (*i.e.,* located in the LUAD-like region; **Figs. 3g-3h**). This result indicates that 3D chromatin organization changes associated with LUAD precede histologic progression in the model.

### Chromatin tracing delineates prognostic and predictive biomarkers in LUAD

To define the relationship between altered 3D genome conformation and gene expression changes during LUAD progression, we performed bulk RNA-seq on dissected GFP+ tumors of *K-MADM-Trp53* mice (**Extended Data Fig. 6d and Supplementary Table 4**). Unsupervised hierarchical clustering revealed two distinct tumor groups (**Extended Data Fig. 6e**) that could be classified as AdenomaG and LUAD cells based on expected changes in gene expression with progression, including loss of lung epithelial markers (AT1/AT2), increased expression of genes connected with gastric identity, epithelial-to-mesenchymal transition (EMT), and metastasis, plus transcriptional alterations associated with enhanced Kras signaling (**Extended Data Figs. 6f-6g**)^40^. We next focused on the scA/B score changes of genes with significantly altered expression between AdenomaG and LUAD cells. Genes with enhanced expression in LUAD cells tended to show increased scA/B scores, whereas those with reduced expression had decreased scA/B scores (**Fig. 4a**). We reasoned that the 3D compartment alterations described above could promote stable gene activation or repression with functional consequences, due to differences in active/repressive epigenetic signatures and chromatin, RNA polymerase II (Pol II), and transcription factor densities in the compartments^41^. Genes with significantly elevated expression in marker genomic loci (**Fig. 4b**) with increased scA/B scores in LUAD cells might be important for LUAD progression, as candidate progression drivers (CPDs) (**Fig. 4c**). In contrast, genes with significantly decreased expression in marker loci with decreased scA/B scores could be candidate tumor suppressors (CTSs) (**Fig. 4c**). We developed a “Trace2Biomarker” pipeline to call CPDs and CTSs (see **Methods**). To test the hypothesis that A/B compartment organization contributes to more stable gene expression regulation, we quantified the gene expression heterogeneity/homogeneity in LUAD cells derived from *K-MADM-Trp53* mice analyzed by single-nucleus RNA-seq (snRNA-seq; **Extended Data Figs. 6h-6j**). Using a published gene expression homogeneity metric^40^, CPD and CTS genes were significantly more homogenously expressed than the top differentially expressed genes (up or down) in regions with unchanged scA/B scores (**Fig. 4d**). We confirmed these findings in a separate single-cell RNA-seq (scRNA-seq) dataset derived from a different *Kras/Trp53* mutant (KP) LUAD mouse model^40^ (**Fig. 4e**). These observations suggest that genes associated with compartment changes during tumor evolution are more homogeneously regulated, leading to more stable regulation of potential cancer driver and suppressor genes than compartment-independent expression regulatory mechanisms.

**Fig. 4.**
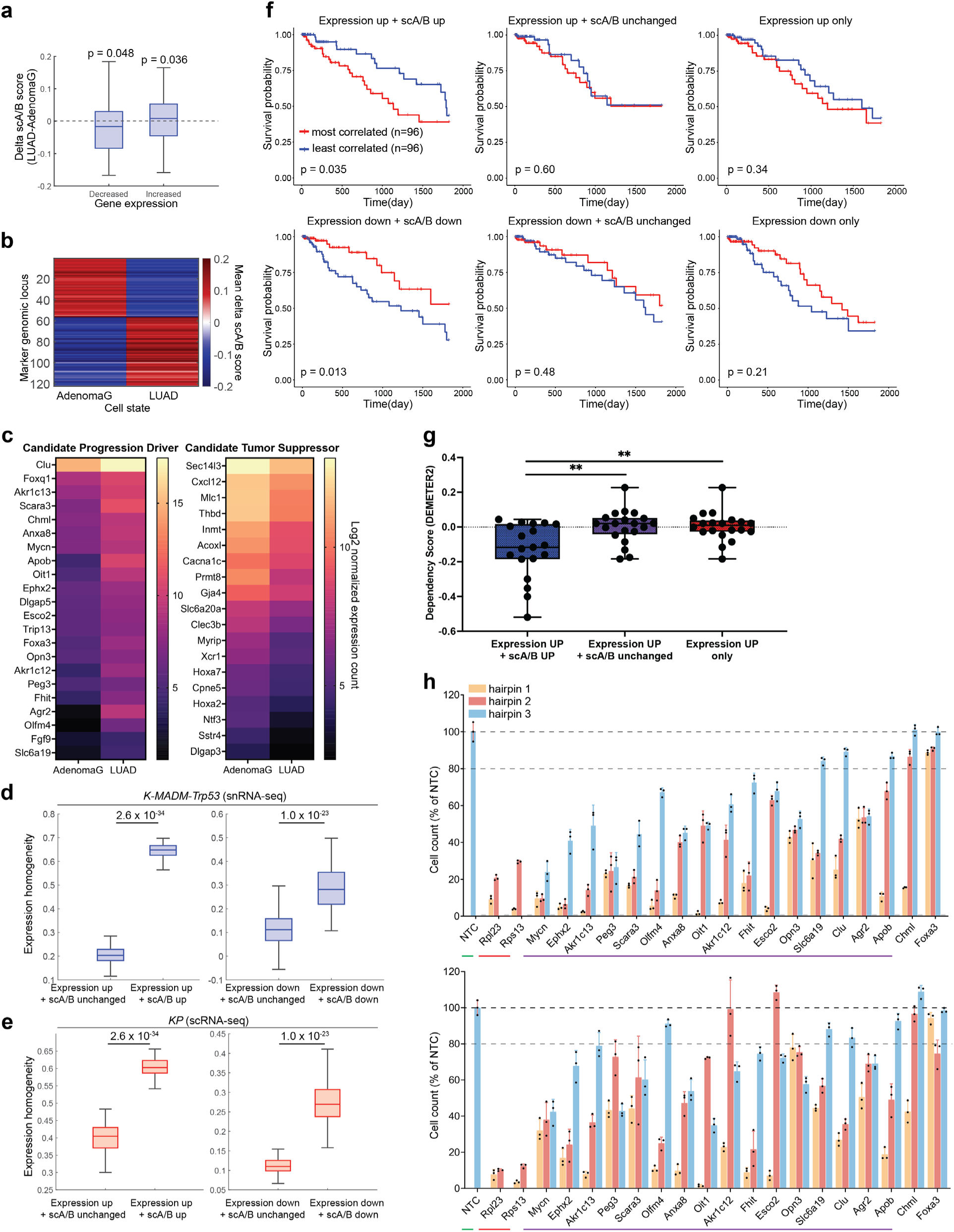
The single-cell 3D genome nominates prognostic genes and genetic dependencies. **a,** scA/B score changes of genes with decreased (n = 94 genes) or increased (n = 120 genes) expression levels from AdenomaG to LUAD cells. p values of one-sided Wilcoxon signed-rank tests are shown. **b,** Mean scA/B score changes in marker genomic loci between AdenomaG and LUAD cells. **c,** Heatmap of candidate progression driver (CPD) and candidate tumor suppressor (CTS) gene expression (log2 normalized expression counts) comparing AdenomaG and LUAD tumors derived from *K-MADM-Trp53* mice. **d-e,** Gene expression homogeneity of LUAD cells in the *K-MADM-Trp53* (single nucleus) (**d**) and KP (single cell) (**e**) model comparing CPDs (Expression up + scA/B up) or CTSs (Expression down + scA/B down) with up/down regulated genes in regions with unchanged scA/B scores. Gene expression homogeneity is quantified with a correlation homogeneity score method^40^. The horizontal lines of each box represent the 75th percentile, median, and 25th percentile. Whiskers extend to the non-outlier maximum and non-outlier minimum. Outliers are defined as values away from the top or bottom of the box by more than 1.5 times interquartile range. **f,** Kaplan-Meier survival curves comparing TCGA LUAD patients with gene expression profiles most or least correlated with CPDs (Expression up + scA/B up), CTSs (Expression down + scA/B down), and the corresponding controls (Expression up + scA/B unchanged or Expression down + scA/B unchanged; Expression up only or Expression down only). Top row: 21 genes are included in each panel. Bottom row: 19 genes are included in each panel. In the control groups, the genes are the ones with the highest expression fold change among all genes fitting the criteria. All analyses were performed with a top vs. bottom 20% (quintiles; n = 96 tumors per group). p values of two-sided log-rank test are shown. **g,** Dependency scores (using the DEMETER2 algorithm) of LUAD cell lines (n = 57) comparing CPDs (Expression up + scA/B up) and corresponding controls (Expression up + scA/B unchanged or Expression up only). n = 19-20 genes per group with available data in RNAi screens in the Cancer Dependency Map. Lower (more negative) score = more dependent. **p < 0.01, Wilcoxon rank-sum test. The horizontal lines of each box represent the 75th percentile, median, and 25th percentile. Whiskers extend to the maximum and minimum. **h,** Cell viability (mean ± SD, normalized to the mean of non-targeting control (NTC), n = 3 replicates per hairpin, 3 hairpins per gene) of arrayed RNAi screen targeting CPD genes in the KP (upper panel) and SA6082inf (lower panel) LUAD cell lines. NTC (green underline), positive controls (red underline), and CPD genes with significant phenotypes (at least two out of three hairpins with less than 80% of NTC cell count, purple underline) in both cell lines are indicated.

To investigate the relative importance of the CPD and CTS genes identified by our analysis, we assessed whether they could serve as useful prognostic (of survival) or predictive (of dependency) biomarkers. Importantly, 5 of the 22 CPD genes and 8 of the 19 CTS genes that we identified have not previously been implicated in lung cancer pathogenesis (**Supplementary Table 5**). To determine prognostic significance, we scored individual patient LUAD samples (from The Cancer Genome Atlas (TCGA)^25^) for their correlation with the CPD and CTS gene signatures. For the CPD gene signature, high-scoring patients (top quintile most correlated tumor gene expression, *n* = 96 patients) exhibited significantly worse survival compared to patients with lower scores (bottom quintile, *n* = 96 patients) whereas the inverse was true for the CTS gene signature (**Fig. 4f**). Comparable analyses of the top differentially expressed genes in genomic regions with unchanged scA/B scores (or irrespective of scA/B score changes), by contrast, were poor predictors of patient survival (**Fig. 4f**). Similarly, CPD genes identified by combining 3D genome organization and gene expression data showed greater gene dependency in LUAD cell lines analyzed in the Cancer Dependency Map^42^ than top differentially expression genes with unchanged scA/B or irrespective of scA/B score (**Fig. 4g**). To directly validate the function of identified CPD genes in LUAD, we performed an arrayed lentiviral RNAi screen targeting 18 CPD genes with readily-available short hairpin RNAs (shRNAs, 3 per gene; **Supplementary Table 6**) in two independent primary LUAD mouse cell lines isolated from the *K-MADM-Trp53* mouse model (SA6082inf) and an analogous *Kras/Trp53* mutant (KP) mouse model^43^. Knockdown of 16 of 18 (89%) tested CPD genes reduced viability in both cell lines (**Fig. 4h**) – including five novel targets in LUAD (*Ephx2*, *Peg3*, *Apob*, *Oit1*, and *Slc6a19*; **Supplementary Table 5**). Only 6.4% of randomly selected genes would have been expected to affect cell proliferation based on analysis of the Cancer Dependency Map^44–47^ (see **Methods**), arguing that combining 3D genome organization and gene expression data offers greater prognostic (of survival) and predictive (of genetic dependency) power than gene expression alone. Our results thus strongly support the utility of 3D genome mapping as an orthologous tissue-based cancer biomarker.

### Fine-scale chromatin tracing reveals enhancer-promoter interactions in candidate progression drivers

For higher resolution assessment of the evolution of local chromatin structure of CPD and CTS genes during lung cancer progression, we took advantage of fine-scale chromatin tracing. We mapped *in situ* nuclear positions of nearly one thousand genomic loci in the *cis*-regulatory regions of 23 genes, including CPD and CTS genes and the known driver oncogenes *Kras* and *Myc* (**Extended Data Fig. 7a**). For each gene, we targeted 40 consecutive 5-kb to 20-kb loci (20 loci for *Foxa3* due to short genomic length) spanning the promoter and candidate enhancers (**Extended Data Fig. 7a**) and collected datasets from a total of 7,511 AT2, Adenoma (R, Y, G), and LUAD cells. The kilobase-resolution chromatin folding became increasingly more compact during AT2 to adenoma to LUAD progression (**Extended Data Fig. 7b**), suggesting that trends in chromatin compaction from adenoma to LUAD depend on scale. To identify enhancer-promoter (E-P) interactions, we accounted for the confounding effects of gene scale compaction and analyzed the normalized inter-loci distances between putative active enhancers and promoters (**Methods, Extended Data Fig. 7c**). We observed that most E-P loops for CPD genes exhibited increased interactions during progression from adenoma to LUAD, concordant with increased gene expression (**Extended Data Figs. 7d and 7e**). In contrast, E-P loops for CTS genes did not show systematic changes (**Extended Data Figs. 7d and 7e**), suggesting that CTS repression during LUAD progression is not regulated at the E-P interaction level and may instead result from global compaction of the gene regions and/or compartment changes. *Kras* showed an increase in E-P loop interactions in LUAD, which may afford an alternative mechanism for enhancing Kras expression in the adenoma-to-LUAD transition beyond amplification^48^ (**Extended Data Figs. 7e and 7f**). Similarly, *Myc* – basal expression of which is required for *Kras*-driven lung tumor initiation^49^ and overexpression of which is sufficient to induce lung tumorigenesis^50^ – showed increased looping between the *Myc* promoter and two putative downstream enhancers in the AT2 to adenoma transition (**Extended Data Figs. 7e and 7f)**. We further found that the *Myc* locus exhibited increased expression and scA/B score changes comparing adenoma to AT2 cells (termed candidate initiation gene; **Supplementary Table 5**), suggesting that both local looping and larger scale compartment alterations may regulate its expression. These data demonstrate the potential to map both fine- and large-scale 3D genome structure with chromatin tracing to clarify associations between chromatin architectures and expression of functionally important genes in cancer.

### Rnf2 regulates 3D genome folding through a non-canonical function

One major goal of our study was to define novel regulators of 3D genome architecture. We first assessed the role of neighboring immune cells in the tumor microenvironment in governing 3D genome organization and found little influence – arguing that 3D genome organization changes during LUAD progression are likely independent of immune cell interactions. Comparing 3D genome organization of AT2/cancer cells during LUAD progression in relation to their spatial proximity to CD45+ cells showed negligible differences in chromatin conformation heterogeneity, chromatin decompaction, chromatin demixing, and A-B compartment polarization (**Extended Data Figs. 8a-8d**). Moreover, in dimensionality reduction analyses, AT2/cancer cells adjacent to and far from immune cells contained similar spectra of single-cell 3D genome conformations (**Extended Data Fig. 8e**).

To identify possible cell-autonomous upstream regulators of cancer 3D genome reorganization, we developed a “Trace2Regulator” pipeline (see **Methods**). The pipeline leveraged a published Binding Analysis for Regulation of Transcription (BART) algorithm^51,52^ to predict transcription factors and chromatin regulators that bind to upregulated genes located in marker loci with increased scA/B scores in the adenoma-to-LUAD transition. We identified Rnf2 as a top candidate. Rnf2 is a major component of the polycomb repressive complex 1 (PRC1) and catalyzes mono-ubiquitination of lysine 119 of histone H2A (H2AK119ub)^53^. Canonically, PRC1 works in concert with polycomb repressive complex 2 (PRC2), which modifies histones by depositing H3K27me3 to repress transcription^54^. We knocked down Rnf2 (shRnf2) in KP LUAD cells (**Fig. 5a**) and saw a significant negative effect on cell viability (**Fig. 5b**). scA/B score changes upon Rnf2 knockdown significantly correlated with scA/B score changes during LUAD progression (**Fig. 5c**), indicating that Rnf2 partially regulates 3D genome reorganization in LUAD cells. Since the predicted binding sites of Rnf2 were associated with increased scA/B scores, we speculated that Rnf2 may preferentially bind A compartment regions. Consistent with this hypothesis, Rnf2 CUT&RUN revealed significantly higher Rnf2 peak density in compartment A topologically associating domains (TADs, also known as contact domains)^55–59^ (**Fig. 5d**).

**Fig. 5.**
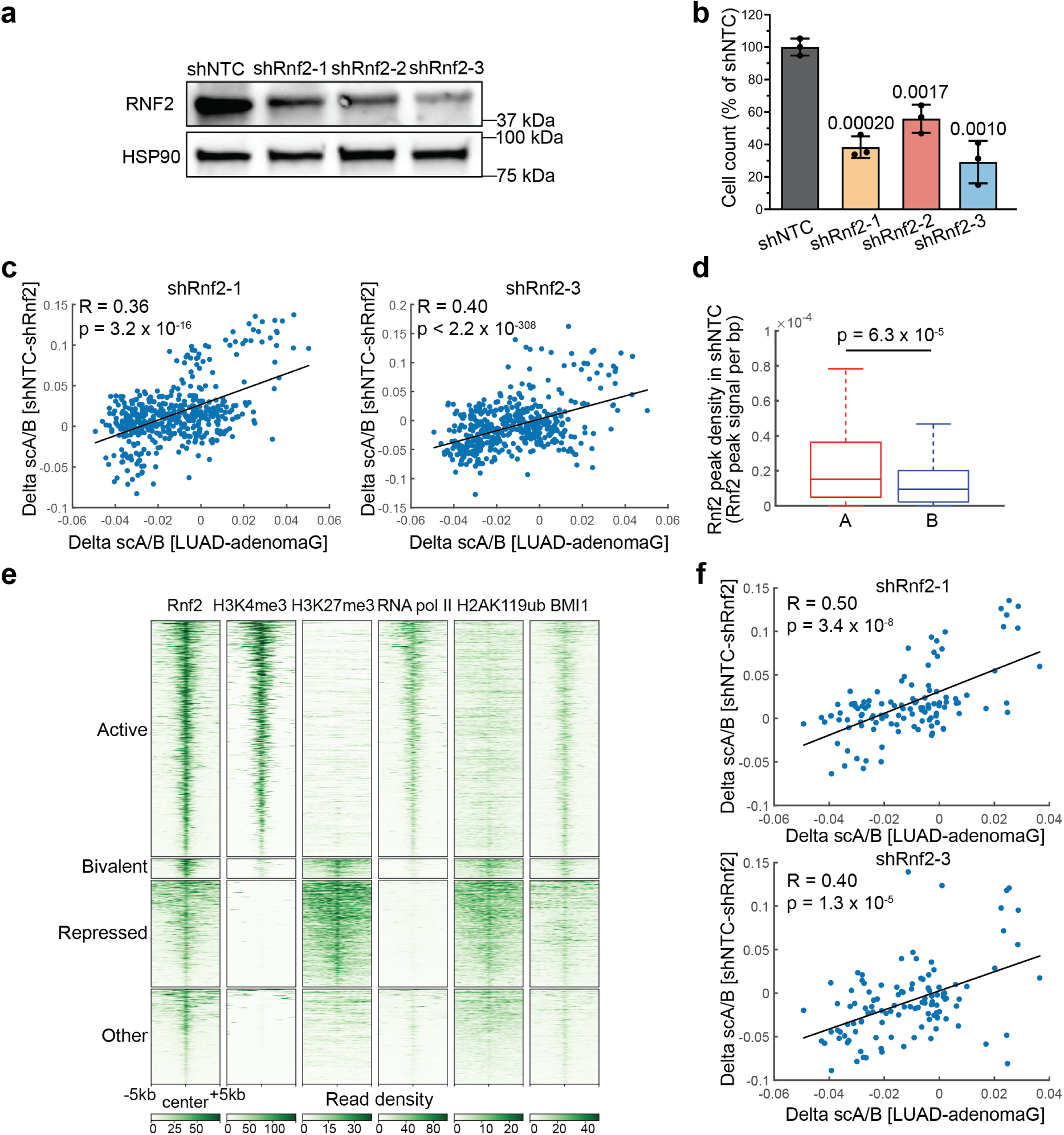
Rnf2 partially regulates 3D genome organization changes during the adenoma-to-LUAD transition. **a,** Western blot analysis of Rnf2 protein levels following Rnf2 knockdown with three independent hairpins in the KP cell line. Control cells were constructed with a non-targeting control shRNA sequence (shNTC). HSP90 is loading control. **b,** Cell viability (mean ± SD, normalized to mean of shNTC) following knockdown with three Rnf2 shRNAs, n = 3 replicates per hairpin. p values from two-sided two-sample t-tests are shown. **c,** scA/B score changes from shRnf2 to shNTC versus those from AdenomaG to LUAD show a significant positive correlation, n = 473 target genomic regions. Black lines are fitted regression lines. Spearman correlation coefficients and p values are shown. **d,** Rnf2 peak densities in A (n = 209 regions) and B (n = 264 regions) compartments in shNTC. The horizontal lines of each box from top to bottom represent the 75th percentile, median, and 25th percentile. Whiskers extend to the non-outlier maximum and non-outlier minimum. Outliers are defined as values at least 1.5 times interquartile range away from the top or bottom of the box. p value of two-sided Wilcoxon rank-sum test is shown. **e,** CUT&RUN read density heatmaps of Rnf2, H3K4me3, H3K27me3, RNA polymerase II with phosphorylated S5 modification, H2AK119ub, and BMI1 in shNTC KP cells. Rnf2 peak regions [−5k, +5k] of all target genomic regions are shown and are categorized as active (H3K4me3+, H3K27me3−), bivalent (H3K4me3+, H3K27me3+), repressed (H3K4me3−, H3K27me3+), or other (H3K4me3−, H3K27me3−) based on chromatin marks. **f,** scA/B score changes from shRnf2 to shNTC versus those from AdenomaG to LUAD show a stronger or similarly positive correlation using target genomic regions with only active Rnf2 peaks, n = 113 target genomic regions.

Although the canonical function of Rnf2 is to mediate gene silencing through polycomb repression^53^, it can also (non-canonically) associate with epigenetically active loci^60,61^. As both polycomb-repressed and active chromatin regions tend to localize in compartment A^62,63^, a key question is whether the Rnf2-controlled regions in LUAD are mainly polycomb-repressed or active chromatin. To answer this question, we aligned and clustered the Rnf2 CUT&RUN peaks in all target genomic regions with H3K4me3 and H3K27me3 CUT&RUN profiles. The majority of Rnf2 peaks (51.6%) resided in active regions (H3K4me3+, H3K27me3−) (**Fig. 5e**), with fewer (23.3% and 4.3%, respectively) being found in repressed (H3K4me3−, H3K27me3+) and bivalent (H3K4me3+, H3K27me3+) regions (**Fig. 5e**). CUT&RUN analysis of RNA Pol II with phosphorylated S5 confirmed the active transcriptional states of the active regions (**Fig. 5e**). In addition, scA/B score changes upon Rnf2 knockdown were better or similarly correlated with the scA/B score changes in LUAD progression, if only active Rnf2-bound regions were included in the correlation analysis (**Fig. 5f**). These results suggest that Rnf2 in active genomic loci may regulate 3D genome reorganization associated with LUAD progression.

To further understand the regulatory role of Rnf2 on the 3D genome, we performed H2AK119ub CUT&RUN, and found that WT Rnf2 peaks in active regions do not colocalize with strong H2AK119ub peaks. In contrast, another core PRC1 component BMI1 colocalizes with Rnf2 in all regions (**Fig. 5e**). These data argue that the canonical ubiquitin ligase function of Rnf2 is dispensable for its role in shaping the 3D genome. To test this hypothesis directly, we performed rescue experiments upon Rnf2 knockdown with vectors expressing wild-type (WT) Rnf2 or a catalytic Rnf2 mutant (I53S) that lacks ubiquitin ligase activity^64^ (**Fig. 6a**). We found that scA/B score changes upon rescue with WT Rnf2 significantly correlated with scA/B score changes during LUAD progression (**Fig. 6b**), validating the specific effect of Rnf2. Strikingly, mutated Rnf2 (I53S) also induced scA/B score changes that mirrored LUAD progression, but with a stronger and more significant correlation than that of WT rescue (**Fig. 6b**). These results thus demonstrate that Rnf2 partially regulates 3D genome changes during the adenoma-to-LUAD progression via a ubiquitin ligase-independent function.

**Fig. 6.**
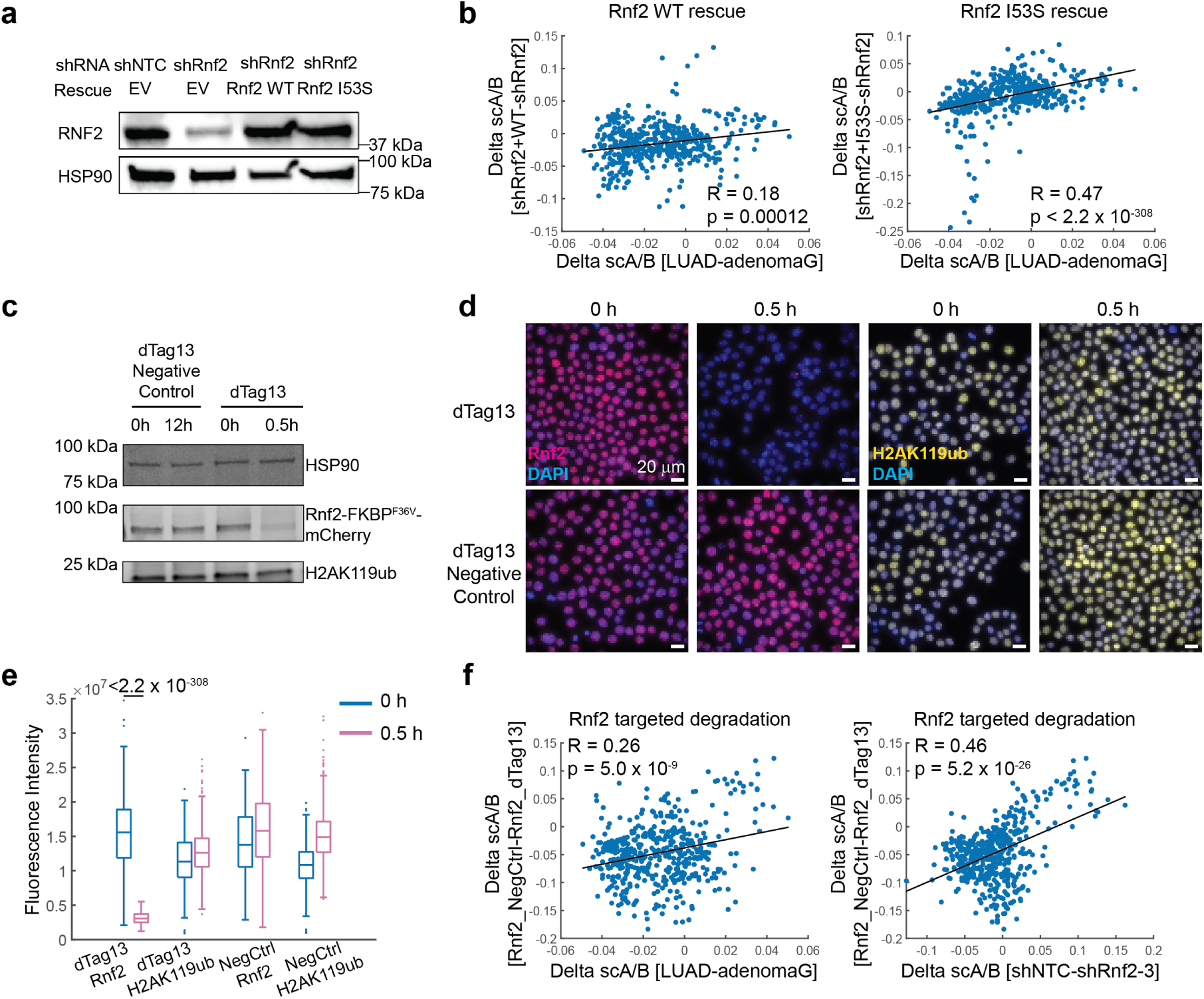
Rnf2 regulates 3D genome organization via a ubiquitin ligase-independent activity. **a,** Western blot analysis of Rnf2 protein levels following rescue of shRNA knockdown (shRnf2-3) with stable transduction of Rnf2 WT, ubiquitin-ligase dead Rnf2 I53S mutant, or empty vector (EV) control. HSP90 is loading control. **b,** scA/B score changes from shRnf2-3 to shRnf2-3+WT Rnf2 (left) and to shRnf2-3+I53S Rnf2 (right) versus those from AdenomaG to LUAD show a stronger positive correlation with expression of a catalytically dead mutant of Rnf2, n = 473 target genomic regions. Black lines are fitted regression lines. Spearman correlation coefficients and p values are shown. **c,** Western blot of Rnf2 and H2AK119ub in KP LUAD cells with Rnf2-dTAG after treatment with the dTAG-13 ligand or negative control ligand (a diastereomer of dTAG-13) at the designated times (0, 0.5, or 12 hours). **d,** Immunofluorescence images of Rnf2 and H2AK119ub in KP LUAD cells with Rnf2-dTAG after treatment with dTAG-13 ligand or negative control at designated times. **e,** Quantification of the immunofluorescence intensities in **d**. The horizontal lines of each box from top to bottom represent the 75th percentile, median, and 25th percentile. Whiskers extend to the non-outlier maximum and non-outlier minimum. Outliers are defined as values at least 1.5 times interquartile range away from the top or bottom of the box. p value from two-sided Wilcoxon rank sum test is shown. **f,** scA/B score changes from Rnf2-degraded cells (dTAG-13) to Rnf2 non-degraded cells (dTAG-13 negative control) versus those from AdenomaG to LUAD and those from shRnf2 to shNTC, with n = 473 target genomic regions. Black lines are fitted regression lines. Spearman correlation coefficients and p values are shown.

Finally, to test whether Rnf2 directly reorganizes the 3D genome in LUAD cells, we used the degradation tag (dTAG) system^65^ to induce acute Rnf2 protein loss in KP cells. We endogenously tagged an FKBP12^F36V^ degron tag to the C-terminus of Rnf2. Rnf2 protein was rapidly degraded after 0.5 h of dTAG ligand (dTAG-13) administration but H2AK119ub levels were unchanged (**Figs. 6c-6e**). We found that the scA/B score changed significantly upon rapid Rnf2 degradation and that these alterations correlated with scA/B score changes during LUAD progression or upon Rnf2 knockdown (**Fig. 6f**). These kinetic degradation experiments support a direct role for Rnf2 in 3D genome reorganization independent of its canonical ubiquitin ligase catalytic function.

## Discussion

In this study, we generated comprehensive 3D genome atlases in physiologically relevant mouse models of lung and pancreatic cancer progression. Beyond establishing a new data rich cancer genomics resource, our analyses support the broad utility of our technologic pipeline to measure and interpret biological heterogeneity of the 3D genome in a variety of biological systems and diseases (**Extended Data Fig. 9**). A key finding from this work is our demonstration that data on single-cell 3D genome organization can distinguish and predict different histologic cancer cell states with high accuracy (**Fig. 3**), despite recent studies showing that 3D chromatin conformation is highly variable even among cells of the same type^66–70^. We further show that high-dimensional 3D genome data can be used to identify genes that govern prognosis and delineate cancer cell dependencies (**Fig. 4**). Perturbation of most of the nominated candidate progression drivers led to defects in cancer cell viability, demonstrating that the single-cell 3D genome organization contains rich information to uncover novel diagnostic and prognostic cancer biomarkers along with genetic vulnerabilities for drug target discovery.

Our results also underscore the potential of high-dimensional *in situ* 3D genome data to identify previously unappreciated 3D genome regulators. By focusing on genes within loci that underwent genome compartment reorganization during LUAD progression, we discovered Rnf2 has a ubiquitin ligase-independent role in reorganizing the 3D genome in active chromatin regions. Although Rnf2 is best known for transcriptional repression^53,54^, recent reports have delineated a non-canonical role for Rnf2 in gene activation, though the precise mechanisms for this function remain controversial^60,61^. In comparison to WT Rnf2, re-expressing a catalytic mutant (I53S) of Rnf2 in LUAD cells more strongly recapitulated 3D genome changes in the adenoma-to-LUAD transition (**Fig. 6b**). One potential explanation for this is that Rnf2 I53S associates more with active chromatin regions than its WT counterpart, facilitating compartment transitions that drive LUAD progression. In support of this model, a recent study demonstrated that H2AK119ub1 and H3K27me3 are required to maintain Rnf2 at canonical PRC2-repressed sites, as loss of Rnf2 catalytic activity displaces it from these sites^71^. An additional layer of regulation may occur downstream of KRAS signaling itself. Prior work in *BRAF*-mutant melanoma^61^ suggested that phosphorylation of Rnf2 by MEK1/2 – a key mediator of the mitogen-activated protein kinase (MAPK) pathway downstream of oncogenic *KRAS* and *BRAF* – allows Rnf2 to recruit activating histone modifiers to poised promoters to induce gene expression. During murine LUAD progression, MAPK signaling downstream of oncogenic *Kras* increases^72,73^, suggesting that a comparable mechanism of Rnf2 activation may be occurring. These data argue that MAPK hyperactivation – permitted by loss of *Trp53*^72,73^ – may facilitate the capacity of Rnf2 to reorganize 3D genome compartmentalization to drive the adenoma-to-LUAD transition. Given increased MAPK signaling in human NSCLC samples^74^, such mechanisms of Rnf2 regulation may also occur during human lung cancer progression.

Recent scRNA-seq^40^ and scATAC-seq^75^ studies in mouse models of *Kras*-driven LUAD characterized the heterogeneous transcriptomic and epigenomic landscapes that arise in cancer, leading to the identification of transitional and plastic cell states with molecular alterations that can drive tumor initiation, progression, and metastasis^20,40,75^. Although transcriptional heterogeneity progressively increased during LUAD progression in these models, we observed a striking non-monotonic evolution of the 3D genome, which was mirrored in PDAC (**Fig. 2**) and could represent a structural bottleneck of the 3D genome in cancer development. In support of a functional role of 3D genome alterations during tumor evolution, combining 3D genome and transcriptomic data led to improved identification of important genes (CPDs and CTSs) that predict prognosis and dependency (**Fig. 4**). One potential mechanism by which nuclear compartment localization may lead to functional importance is by maintaining more stable gene activation or repression due to epigenetic state, chromatin density, proximity to activating/repressive nuclear landmarks, and local interacting protein density (including RNA Pol II and transcription factors)^41^. Consistent with this hypothesis, we observed that differentially expressed genes that resided in loci that underwent concordant compartment transitions were more homogeneously expressed or repressed in LUAD (**Figs. 4d and 4e**). To directly map 3D genome alterations to transcriptionally and epigenetically defined cell states, future studies should combine genome-wide chromatin tracing with imaging-based spatial transcriptomics^76–79^, epigenomics^80^, and proteomics^68,81^ in the same single cancer cells within the native tumor microenvironment. In addition, genome-wide finer-scale chromatin tracing with kilobase resolution may systematically reveal gene-specific enhancer-promoter interactions further defining the molecular features and plasticity of cells during LUAD progression.

Chromatin tracing offers several unique advantages among single-cell 3D genomics methods^35,82–87^ in studying the cancer 3D genome. First, as an imaging-based approach, it retains the native tissue architecture *in situ*, which is essential for pathologic analyses in cancer studies and allows analyses of cell-cell interactions in shaping the 3D genome within the tumor microenvironment (**Extended Data Fig. 8**). Second, the chromatin tracing approach directly traces the 3D folding path of chromatin, whereas sequencing-based approaches indirectly infer 3D chromatin folding based on contact events. Third, the chromatin tracing method allows integration with multiplexed fluorescent protein/immunofluorescence imaging in the same single cells, enabling cell type-specific identification (*e.g.*, SPC+ AT2 cells, CK19+ duct cells) and lineage labels (*e.g.*, MADM), whereas the potential for similar multiplexing is limited in sequencing-based methods. Fourth, the chromatin tracing approach achieves higher cell throughput due to its cost efficiency. For example, a recent scSPRITE study^85^ profiled 1,000 cells at 40-kb to 1-Mb resolution, while our work analyzed over 61,000 cells at 5-kb to 5-Mb resolution, albeit with lower genomic coverage intrinsic to the targeted FISH approach.

Our study demonstrates proof-of-principle of the utility of *in situ* chromatin tracing to garner insights into tumor biology, taking advantage of the capacity to visualize the entire spectrum of lung and pancreatic tumorigenesis in mouse models with genetically encoded lineage markers for tracing subclonal evolution. Furthermore, the choice of model system allowed us to account for differences in cell cycle progression and copy number variation (CNV) across histologic states, which could confound 3D genome analyses. We previously showed with pulse 5-ethynyl-2’-deoxyuridine (EdU) incorporation experiments in *K-MADM-Trp53* mice that ∼2% of preinvasive adenoma cells are actively proliferating compared to ∼8% of LUAD cells^30^. Therefore, most of our profiled cancer cells are in G_0_/G_1_, which should have minimal confounding effects on chromatin folding and nuclear organization^88^. Furthermore, the monotonic increase in the proportion of cells proliferating in going from the AT2 (post-mitotic) to adenoma to LUAD populations cannot explain the reversible trend of 3D genome changes observed across these states. Whole-exome sequencing of AdenomaG and LUAD cells isolated from *K-MADM-Trp53* mice showed only a subset of LUAD tumors exhibiting evidence of gains of Chromosome 6 and losses of Chromosomes 4, 9, and 11 (**Extended Data Figs. 6a-6b**), as has been observed in other *Kras/Trp53* mutant murine LUAD models^40,48^. Given the lack of systematic gains or losses across most chromosomes and tumors, it is unlikely that CNVs could account for the global chromatin structure differences observed between adenoma and LUAD cells across all autosomes. Finally, few single nucleotide variants (SNVs) were observed in both adenoma and LUAD tumors, none of which resulted in protein-altering mutations in known cancer driver genes^89^ (**Extended Data Fig. 6c and Supplementary Table 7**), similar to findings in other genetically engineered mouse LUAD models^48,90,91^. These data argue that the observed 3D genome alterations are not merely a surrogate of the mutational evolution of cancer.

In contrast to the mouse, human tumors exhibit significantly more complex somatic genotypes compared with those in genetically engineered mouse models of a similar type^48^. Indeed, the human NSCLC genome is highly heterogeneous, with different tumors displaying distinct combinations of point mutations, structural variations, and fusion genes^92^. It therefore remains to be explored whether the cell state-specific 3D genome architectures observed in murine LUAD progression are recapitulated in human lung tumors, and how 3D genome evolution differs across driver oncogenes (e.g., *KRAS* vs. *EGFR*). To answer these questions, one can apply genome-wide chromatin tracing to rare clinical human lung tumor samples harboring a preinvasive component (carcinoma *in situ*) adjacent to invasive tumor regions and normal regions. As DNA FISH has been traditionally used to map structural variations in cancer, the FISH-based chromatin tracing technique has the potential to delineate multi-dimensional genomic alterations in single cancer cells, which may offer even more robust biomarkers for cancer diagnosis and therapeutics. Given the rich information one can derive from mapping genome-wide true-3D chromatin folding in subclonal cancer progression, we anticipate that comprehensive tissue-based chromatin tracing – when applied broadly in cancer and combined with multi-modal spatial profiling of the transcriptome, epigenome, proteome, and metabolome – will have transformative potential to reveal new aspects of cancer biology and novel biomarkers for cancer diagnosis, subtype stratification, prediction of treatment response, and therapeutic development.

## Methods

### Mice

#### Mouse strains

Animal studies were approved by the Institutional Animal Care and Use Committee of Yale University. Mice were housed in a specific-pathogen free facility with controlled temperature and day/night cycles and maintained in a mixed background. *MADM11-GT* (Stock #013749), *MADM11-TG* (Stock #013751), and *Pdx1-Cre* (Stock #014647) were obtained from the Jackson Laboratory (JAX). *LSL-Kras^G12D^* (JAX Stock #008179) and *Trp53^KO^*(JAX Stock #002101) mice were a gift from Dr. Tyler Jacks. *LSL-Kras^G12D^/Kras^WT^; MADM11-TG,Trp53^KO^/MADM11-TG,Trp53^WT^* breeder mice were generated as previously described^30^. *LSL-Kras^G12D^/Kras^WT^; MADM11-TG, Trp53^KO^/MADM11-TG, Trp53^WT^* were crossed with *MADM11-GT* mice (with or without *Pdx1-Cre*) to generate *K-MADM-Trp53* or *Pdx1-Cre; K-MADM-Trp53* experimental mice of both sexes for lung and pancreatic cancer analyses, respectively. Genotyping primers and protocols have been previously reported^30^.

#### Lentivirus production and infection

pPGK-Cre lentiviral backbone was amplified and transfected into 293T cells using VSV-G (Addgene 8454) and psPAX2 (Addgene 12260) packaging vectors and TransIT-LT1 kit (MirusBio). After 48 and 72 hours, virus was collected, filtered, and ultracentrifuged. Lentivirus was resuspended in OptiMEM (ThermoFisher) and administered intratracheally to *K-MADM-Trp53* mice at 6-10 weeks of age to generate lung tumors.

### Probe design and synthesis

#### Template probe design

In genome-wide chromatin tracing, to select target genomic regions for mChr1-mChr19, we first downloaded TAD coordinates in mouse embryonic stem cells from http://3dgenome.fsm.northwestern.edu/download.html. Second, we selected a list of target TADs containing one or more of the following features: (1) TADs containing MAPK pathway genes; (2) TADs containing “classic” oncogenes and tumor suppressor genes; and (3) TADs containing super-enhancers. Third, to select other target TADs, we selected 30 TADs that are equally spaced along each entire chromosome and removed those that are within an interval to the feature-containing target TADs. We then combined the remaining other target TADs and feature-containing target TADs as the target genomic regions. We designed about 400 oligos targeting a 60-kb feature-containing region within each feature-containing TAD, and about 400 oligos targeting the central 60-kb region of other target TADs. A total of 50 target TADs were designed for mChr6 and 18-25 target TADs were designed for each of the other autosomes. A total of 473 target TADs spanning mChr1-mChr19 were selected. To distinguish the identities of each target TAD, we adapted a previously published Hamming weight 2 (HW2) binary barcode design strategy^18^. We rearranged the binary code assignment to (1) minimize the variance of the number of TADs imaged in each imaging round across different imaging rounds for each chromosome to make sure no more than 1 TAD was imaged per imaging round for each chromosome, except for mChr6; and (2) for mChr6, maximize the genomic distance between the TADs imaged in the same imaging round. The template oligos for the 473 target TADs were designed as follows: each template oligo consisted of 6 regions (from 5’ to 3’): (1) a 20-nucleotide (20-nt) forward priming region; (2) a 20-nt adapter binding region; (3) a 40-nt genome targeting region; (4) a second 20-nt adapter binding region; (5) a third 20-nt adapter binding region; and (6) a 20-nt reverse priming region. The forward and reverse 20-nt priming sequences were generated from random sequences that were screened to lack homology to the mouse genome and optimized for PCR amplifications. The 20-nt adapter binding sequences bind to 60-nt adapter oligos consisting of one 20-nt adapter sequence and two 20-nt readout probe binding sequences. Both the adapter and readout probe binding sequences were generated from random sequences with minimum homology to the mouse genome and maximum performance in signal to noise ratios. The 40-nt genome targeting sequences were designed with OligoArray2.1^93^ using the following parameters: (1) the melting temperatures of the target sequences were between 65 °C and 85 °C; (2) the melting temperatures of potential secondary structures were less than 76 °C; (3) the melting temperatures of potential cross-hybridizations were less than 72 °C; 4) the GC content of the sequences was between 20% and 90%; (5) no consecutive repeats of six or more A’s, T’s, C’s and G’s were identified; and (6) adjacent oligos were allowed to have 30-nt overlapping sequences. Genome targeting sequences were then screened against the mouse genome with BLAST+ to ensure single matches to the reference genome^94^. Genomic targeting sequences were further screened against a mouse repetitive database from Repbase (https://www.girinst.org/repbase/)^95^. Sequences with more than 16-nt homology to a list of repetitive sequences in the mouse repetitive database were removed. A monolayer of 400 oligos were then selected for each target TAD to construct the template probe library. The template probe library pool was purchased from Twist Biosciences. Template probe sequences are provided in **Supplementary Table 1**. Genome coordinates (mm9), codebook and target TAD features are provided in **Supplementary Table 2**.

In fine-scale chromatin tracing, we designed probes that target the regulatory regions of *Kras*, *Myc*, 13 CPD genes and 8 CTS genes. Genomic regions of interest were chosen based on gene annotations and previously published Hi-C data in mES cells^96^. Putative enhancers were identified by the union of Ensembl predicted enhancers and H3K4me1 (ENCFF536DWZ) and DNaseI (ENCFF268DLZ) intersected ChIP-seq peaks in adult mouse lung. We selected 40 consecutive 5-kb to 20-kb genomic loci (20 loci for *Foxa3* due to shorter genomic length) for each gene. We performed 50 rounds of three-color imaging to decode the identity of each locus. For the first 40 rounds, each 5-kb to 20-kb genomic locus was read out one at a time on all genes (seven genes in 560-nm channel, eight genes in 647-nm channel, eight genes in 750-nm channel) in each hybridization round. The last 10 rounds read out one gene at a time in the three channels. We designed 150 oligos per 5kb genomic region. The template oligos were designed as follows: each template oligo consisted of 6 regions (from 5’ to 3’): (1) a 20-nucleotide (20-nt) forward priming region; (2) a 20-nt genomic loci readout region; (3) a 30-nt genome targeting region; (4) a second 20-nt genomic loci readout region; (5) a 20-nt gene readout region; and (6) a 20-nt reverse priming region. The forward and reverse 20-nt priming sequences and readout sequences were generated as described above. The 30-nt genome targeting sequences were designed with ProbeDealer^97^ using the following parameters: (1) the melting temperatures of the target sequences were between 66 °C and 100 °C; (2) the melting temperatures of potential secondary structures were less than 76 °C; (3) the melting temperatures of potential cross-hybridizations were less than 72 °C; 4) the GC content of the sequences was between 30% and 90%; (5) no consecutive repeats of six or more A’s, T’s, C’s and G’s were identified; and (6) adjacent oligos were allowed to have 20-nt overlapping sequences. Genome targeting sequences were then screened against the mouse genome with BLAST+ to ensure single matches to the reference genome^94^. Genomic targeting sequences were further screened against a mouse repetitive database from Repbase (https://www.girinst.org/repbase/)^95^. Sequences with more than 16-nt homology to a list of repetitive sequences in the mouse repetitive database were removed. A monolayer of 150 oligos per 5-kb genomic region was then selected for each target locus to construct the template probe library. The template probe library pool was purchased from Agilent. Template probe sequences are provided in **Supplementary Table 1**. Genome coordinates (mm10) are provided in **Supplementary Table 2**.

#### Primary probe synthesis

Primary probes were synthesized from the template probe library using previously published protocols following a procedure of limited-cycle PCR, *in vitro* transcription, reverse transcription, and probe purification^11,77,98,99^. All PCR primers, reverse transcription primers, adapters, and readout probes were purchased from Integrated DNA Technologies (IDT). The sequences are provided in **Supplementary Table 3**.

### Mouse *K-MADM-Trp53* lung tissue chromatin tracing experiments

#### Tissue preparation

*K-MADM-Trp53* mice were dissected at sign of respiratory distress via CO_2_ asphyxiation. Lungs were perfused with ice cold 1×PBS, incubated in 4% (vol/vol) paraformaldehyde (Electron Microscopy Sciences, 15710-S) overnight, and cryoprotected with 30% sucrose (Sigma-Aldrich, S0389). Tissue was embedded in Tissue-Tek OCT (Sakura, 4583) and frozen on dry ice before storage at -80 °C.

#### Coverslip treatment

Prior to tissue sectioning, coverslips (Bioptechs, 40-mm-diameter, #1.5) were first silanized as previously described^100,101^. In brief, coverslips were immersed into a 1:1 mixture of 37% (vol/vol) hydrochloric acid (HCl) and methanol at room temperature for 30 min. Coverslips were then washed with deionized water three times, followed by a 70% ethanol wash. Coverslips were dried in a 70 °C oven for 1 h and immersed into chloroform containing 0.2% (vol/vol) allyltrichlorosilane (Sigma, 107778) and 0.1% (vol/vol) triethylamine (Millipore, TX1200) for 30 min at room temperature. Coverslips were then washed with chloroform and ethanol and dried in a 70 °C oven for 1 h. Silanized coverslips can be stored in a desiccated chamber at room temperature for weeks. For tissue attachment, silanized coverslips were treated with 1% (vol/vol) polyethylenimine (Sigma, 408727) in water for 5 min and washed twice in water. Coverslips were then air-dried and ready for tissue attachment.

#### Tissue sectioning

Frozen mouse lung tissue blocks were sectioned at a thickness of 10 μm at - 20°C on a cryostat. Three consecutive tissue slices were sectioned at a time: one for hematoxylin and eosin (H&E) staining, one for whole-section fluorescence imaging, and one for chromatin tracing. The sections were air-dried at room temperature for 1 h and then directly used or stored at -20°C for months.

#### H&E staining and whole-section fluorescence imaging

H&E staining was performed by Yale Pathology Tissue Services (YPTS). For whole-section fluorescence imaging, the section was mounted in VECTASHIELD Vibrance Antifade Mounting Medium with DAPI (VectorLabs, H-1800-2). Stitched fluorescence images in 353-nm, 488-nm, and 592-nm illumination channels for DAPI, GFP, and tdTomato, respectively, were collected with a Plan-Apochromat 10×/0.45 M27 objective on a Zeiss Axio Imager M2 microscope.

#### Co-immunofluorescence and fluorescent protein imaging of the chromatin tracing section

Frozen tissue sections were first balanced at room temperature for 10 min. Tissue sections were then hydrated with DPBS (Gibco, 14190-144) for 5 min twice. Tissue sections were permeabilized with 0.5% (vol/vol) Triton X-100 (Sigma-Aldrich, T8787) in DPBS for 30 min at room temperature and washed in DPBS for 2 min twice. Tissue sections were then blocked for 30 min at room temperature in blocking buffer containing 1% (wt/vol) BSA (Sigma-Aldrich, A9647-100G), 22.52 mg/mL glycine (AmericanBio, AB00730), 10% (vol/vol) donkey serum (Millipore Sigma, S30-100ml), 5% (vol/vol) goat serum (Invitrogen, 31873) and 0.1% (vol/vol) Tween-20 (Sigma-Aldrich, P7949) in DPBS. Tissue sections were incubated with rabbit anti-SPC antibody (Millipore, AB3786, 1:50) and rat anti-CD45 antibody (BioLegend, 103101, 1:100) in blocking buffer at 4 °C overnight. The samples were washed with DPBS for 5 min three times and then incubated with DyLight 800-labeled donkey anti-rabbit secondary antibody (Invitrogen, SA5-10044, 1:1000) and Alexa Fluor 647-labeled goat anti-rat secondary antibody (Invitrogen, A-21247, 1:1000) in blocking buffer for 1 h at room temperature. The samples were washed in DPBS for 5 min three times and then incubated with DAPI (Thermo Fisher, 62248) at 1:1000 dilution in 2× SSC (diluted from 20× SSC, Invitrogen, 15557-044) for 10 min. The samples were then mounted onto a Bioptechs FCS2 flow chamber and replenished with imaging buffer with an oxygen scavenging system (50 mM Tris-HCl pH 8.0 (AmericanBio, AB14043), 10% wt/vol glucose (Sigma-Aldrich, G5767), 2 mM Trolox (Sigma-Aldrich, 238813), 0.5 mg/mL glucose oxidase (Sigma-Aldrich, G2133), 40 μg/mL catalase (Sigma-Aldrich, C30)). The imaging buffer was freshly prepared for each experiment and was covered with a layer of mineral oil (Sigma, 330779) in the reservoir tube to prevent continuous oxidation. We then selected multiple fields of view (FOVs) at predefined tumor regions based on tumor grades of the adjacent whole-section fluorescence images. Tumor grades were confirmed independently by two investigators (S.S.A. and M.D.M.). At each FOV, we sequentially took z-stack images of DAPI, GFP fluorescence, tdTomato fluorescence, anti-SPC immunofluorescence and anti-CD45 immunofluorescence with 405-nm, 488-nm, 560-nm, 647-nm, and 750-nm laser illuminations. The z-stacks (7-10 μm total) had a step size of 200-nm and an exposure time of 0.4 s at each step.

#### Primary probe hybridization

After imaging, tissue sections on coverslips were de-assembled from the chamber, treated with 1 μg/mL proteinase K (Invitrogen, AM2546) in 2% (vol/vol) SDS (Sigma-Aldrich, 05030-1L-F) in 2× SSC at 37 °C for 10 min, and washed in DPBS for 2 min twice. Tissue sections were then treated in 0.1 M HCl for 5 min and washed in DPBS for 2 min twice. Tissue sections were digested with 0.1 mg/mL RNase A (ThermoFisher Scientific, EN0531) in DPBS for 45 min at 37 °C and washed with 2× SSC for 2 min twice. Tissue sections were treated with pre-hybridization buffer containing 50% (vol/vol) formamide (Sigma-Aldrich, F7503) and 0.1% (vol/vol) Tween-20 in 2× SSC. Synthesized primary probes were dissolved in 25 μL hybridization buffer containing 50% (vol/vol) formamide and 20% (vol/vol) dextran sulfate (Millipore, S4030) in 2× SSC. The total probe concentration was 30-40 μM. Coverslips were immersed in hybridization buffer containing probes in 60 mm petri dishes, heat denatured in a 90 °C water bath for 3.5 min, and subsequently incubated the petri dish at 47 °C in a humid chamber for 36-48 hours. Samples were washed with 50% (vol/vol) formamide in 2× SSC for 15 min twice at room temperature followed by 2× SSC for an additional 15 min at room temperature. Washed samples were then incubated with 0.1-μm yellow-green beads (Invitrogen, F8803) resuspended in 2× SSC as fiducial markers for drift correction, washed with 2× SSC briefly, incubated with DAPI at 1:1000 dilution in 2× SSC for 10 min for image registration, and washed again with 2× SSC for 2 min twice.

#### Readout probe hybridization and imaging

After the primary probe hybridization, the sample was repeatedly hybridized with adapters and readout oligonucleotide probes, imaged, and photobleached for a total of 50 rounds. Each adapter probe was 60-nt and consisted of a 20-nt primary probe binding region and two consecutive 20-nt readout probe binding regions. The readout probes were 30-nt oligos conjugated with Alexa Fluor 647 (or Cy5) or ATTO 565 (or Cy3) fluorophores with 20-nt complementary to the readout probe binding regions of adapters. The sequences of adapter and readout probes are provided in **Supplementary Table 3**. To perform automatic buffer exchange during multiple rounds of readout probe hybridization and imaging, we used a Bioptechs FCS2 flow chamber and a computer-controlled, custom-built fluidics system^11,77^. Prior to sequential readout probe hybridization and imaging, we used DAPI images for registration to images taken before primary probe hybridization. We assembled the sample onto the Bioptechs FCS2 flow chamber and further flowed 2 mL imaging buffer through the chamber. We selected the same FOVs as those selected during co-immunofluorescence and fluorescent protein imaging. At each FOV, we took z-stack images with 488-nm and 405-nm laser illuminations to image fiducial beads and DAPI respectively. The z-stacks (7-10 μm total) had a step size of 200-nm and an exposure time of 0.4 s at each step. For each round of readout probe hybridization, we flowed through the chamber 2 mL readout probe hybridization buffer (20% vol/vol ethylene carbonate (Sigma-Aldrich, E26258) in 2× SSC) containing two adapter probes each at 50 nM concentration and incubated for 15 min at room temperature. We flowed through the chamber 2 mL wash buffer (20% vol/vol ethylene carbonate in 2× SSC) for 2 min and further flowed through 2 mL readout probe hybridization buffer containing the two dye-labeled readout probes each at 75 nM concentration. We incubated the sample for 15 min at room temperature and further flowed through 2 mL wash buffer for 2 min. We then flowed 2 mL imaging buffer through the chamber. At each FOV, we took z-stack images with 647-nm, 560-nm, and 488-nm laser illuminations. Dye-labeled readout probes were imaged in the 647-nm and 560-nm channels and fiducial beads were imaged in the 488-nm channel. The z-stack (7-10 μm total) had a step size of 200-nm and an exposure time of 0.4 s at each step. After imaging, we switched buffer to readout probe hybridization buffer containing 1 μM dye-free readout probes (blocking oligos) and photobleached the sample by continuous simultaneous laser illuminations with 750-nm, 647-nm, and 560-nm lasers for 40 s. We then flowed 5 mL 2× SSC through the chamber to wash away the unbound dye-free readout probes before the next hybridization round. A total of 50 hybridization rounds was performed. The color shift between the 647-nm and 560-nm channels was canceled by taking z-stack calibration images of 100-nm Tetraspeck beads (Invitrogen, T7279) attached to a coverslip surface.

### Mouse *K-MADM-Trp53* lung tissue fine-scale chromatin tracing

We performed fine-scale chromatin tracing targeting in total 1200 genomic loci in the *cis* regulatory regions of 30 genes, including 14 candidate progression drivers (CPDs), 14 candidate tumor suppressors (CTSs), *Kras*, and *Myc*. For each gene, we target 40 (20 for *Foxa3* due to short gene length) consecutive 5-kb to 20-kb loci spanning the promoter and candidate enhancers. To distinguish the identity of each target locus in each gene region, we used a parallel chromatin tracing approach: each template probe consists of (from 5’ to 3’) a 20-nt forward priming region, a 20-nt locus-specific readout sequence, a 30-nt genome-targeting sequence, a second 20-nt locus-specific readout sequence, a 20-nt gene-specific readout sequence and a 20-nt reverse priming sequence. We divided the 30 genes into three fluorescence channels (560-nm, 647-nm, 750-nm), with 10 genes profiled per channel. In each channel, probes targeting each of the 40 loci shared the same locus-specific readout sequence. Probes targeting the same gene region shared the same gene-specific readout sequence. The template probe library pool was purchased from Agilent (**Supplementary Table 1**). Primary probes were synthesized following the protocols described above. We first hybridized primary probes to the genome. The hybridization procedure for the fine-scale chromatin tracing is similar to that of the megabase-resolution chromatin tracing except that (1) heat denaturation was performed by incubating the petri dish in a 90 °C water bath for 4 min; and (2) primary probes were incubated in a petri dish at 40 °C in a humid chamber for 36-48 hours. We then sequentially hybridized dye-labeled readout probes and performed 50 rounds of three-color imaging, including 40 rounds for locus-specific readout and 10 rounds for gene-specific readout. The sequences of adapter and readout probes are provided in **Supplementary Table 3**.

### Mouse *K-MADM-Trp53* pancreas tissue chromatin tracing experiments

The experimental procedure for the pancreas tissue is similar to that of the lung tissue except that pancreas tissue sections were permeabilized with 0.5% (vol/vol) Triton X-100 (Sigma-Aldrich, T8787) in DPBS for 20 min at room temperature, washed in DPBS for 2 min twice, incubated with rabbit anti-Cytokeratin 19 antibody (Abcam, ab52625, 1:50 and rat anti-Ki-67 antibody (Invitrogen, 14-5698-82, 1:100) in blocking buffer at 4 °C overnight. Heat denaturation was performed by incubating the petri dish in a 90 °C water bath for 3 min.

### Mouse cell line chromatin tracing experiments

#### Cell culture

The KP mouse primary adenocarcinoma cell line, 31671, was a gift from Dr. Nik Joshi and was derived from an autochthonous *LSL-Kras^G12D^/Kras^WT^; p53^flox/flox^* mouse administered with Adeno-Cre^102^. The *K-MADM-Trp53* mouse primary adenocarcinoma cell line, SA6082inf, was derived from collagenase-based dissociation of a large green tumor dissected from a *K-MADM-Trp53* mouse. *Kras* and *Trp53* genotypes were confirmed by PCR. Both cell lines were cultured in DMEM (Corning, 10-013-CV) containing 10% (vol/vol) FBS (Gibco, 26140-079) and 1% (vol/vol) Penicillin-Streptomycin (ThermoFisher, 15140-122) at 37 °C with 5% CO_2_. Cells were passaged whenever they reached confluency. Both cell lines were seeded onto UV-sterilized coverslips (Bioptechs, 40-mm-diameter, #1.5) and permitted to grow until 60-70% confluency prior to primary probe hybridization. All cell lines were determined to be free of mycoplasma via PCR (ATCC, 30-1012K).

#### Stable Rnf2 knockdown and rescue cell line construction

Lentiviral supernatant for Sigma Mission^TM^ shRNAs targeting *Rnf2* and a non-targeting control (shNTC) was obtained from the Yale Cancer Center Functional Genomics Core. KP cells were seeded in 6-well plates (Falcon, 353046) with a density of 50,000 cells/well. Cells were infected with lentivirus at different titers 24 hours after seeding. Stable transfected cells were selected with 6 μg/mL puromycin (Gibco, A11138-03) for 48-72 hours after lentiviral transfection. Cell pellets were collected after 48 hours of selection for quantitative RT-PCR validation and after 72 hours of selection for western blot validation. For Rnf2 rescue experiments, Rnf2 WT and Rnf2 I53S cDNAs (both harboring a mutated Rnf2 shRNA target seed sequence) were synthesized from Genscript and inserted into the multiple cloning site of pLV-EF1a-IRES-Hygro (Addgene 85134). To construct rescue cell lines (shNTC+empty vector, shRnf2+empty vector, shRnf2+I53S Rnf2), stable Rnf2 knockdown KP cells were seeded in 6-well plates (Falcon, 353046) with a density of 50,000 cells/well. Cells were infected with lentivirus at different titers 24 hours after seeding. Stable transfected cells were selected with 700 μg/mL hygromycin (Gibco, 10687010) for 48-72 hours after lentiviral transduction. Rnf2 protein expression was confirmed by western blot.

#### Primary probe hybridization

Cells were washed with DPBS twice for 2 min each, fixed with 4% (vol/vol) paraformaldehyde (EMS, 15710) in DPBS for 10 min, and washed twice with DPBS for 2 min each. Cells were permeabilized with 0.5% (vol/vol) Triton-X (Sigma-Aldrich, T8787) in DPBS for 10 min and washed twice with DPBS for 2 min each. Next, the cells were treated with 0.1 M HCl for 5 min at room temperature, washed with DPBS twice for 2 min each, treated with 0.1 mg/mL RNase A in DPBS for 45 min at 37 °C, and washed twice with 2× SSC for 2 min each.

The cells were subsequently incubated in pre-hybridization buffer containing 50% (vol/vol) formamide and 0.1% (vol/vol) Tween-20 in 2× SSC. Synthesized primary probes were dissolved in 25 μL hybridization buffer containing 50% (vol/vol) formamide and 20% (vol/vol) dextran sulfate in 2× SSC. The final probe concentration was 30-40 μM. We then added the hybridization buffer containing probes to a 60 mm petri dish and flipped the coverslip onto it so that the cells were immersed into hybridization buffer. Heat denaturation was performed by incubating the petri dish in a 90 °C water bath for 4 min. The petri dish was incubated at 47 °C in a humid chamber for 36-48 hours. After hybridization, the cells were washed with 50% (vol/vol) formamide in 2× SSC for 15 min twice at room temperature and washed with 2× SSC for an additional 15 min. We then incubated each sample with 0.22-μm light yellow beads (Spherotech, FP-0245-2) resuspended in 2× SSC as fiducial markers for drift correction and washed the sample with 2× SSC briefly.

#### Readout probe hybridization and imaging

We followed the same “Readout probe hybridization and imaging” procedure, as performed in “*Mouse K-MADM-Trp53 lung tissue chromatin tracing experiments*” described above, except that (1) DAPI images were acquired after sequential readout probe hybridization and imaging; (2) both readout probe hybridization buffer and wash buffer were composed of 35% (vol/vol) formamide in 2× SSC instead of 20% (vol/vol) ethylene carbonate; (3) fiducial beads were imaged in the 405-nm channel; and (4) tris(2-carbox-yethyl)phosphine (TCEP; Sigma-Aldrich, C4706) cleavage was used instead of photobleaching for fluorescence signal removal between the readout hybridization rounds. TCEP reduces disulfide bonds connecting fluorophores to readout probes, removing fluorophores from readout probes. The TCEP cleavage buffer was composed of 50 mM TCEP and 1 μM dye-free readout probes in 20% (vol/vol) ethylene carbonate to block unoccupied readout probe binding regions on adapters from interfering with the next round of hybridization and imaging.

### Microscope setup

For imaging, we used a custom-built microscope with a Nikon Ti2-U body, a Nikon CFI Plan Apo Lambda 60× Oil (NA1.40) objective lens, and an active auto-focusing system^103^. Different laser settings were applied for imaging tissues or cell lines. For *K-MADM-Trp53* lung chromatin tracing experiments, illumination lasers included: a 750-nm laser (2RU-VFL-P-500-750-B1R, MPB Communications), a 647-nm laser (2RU-VFL-P-1000-647-B1R, MPB Communications), a 560-nm laser (2RU-VFL-P-1000-560-B1R, MPB Communications), a 488-nm laser (2RU-VFL-P-500-488-B1R, MPB Communications), and a 405-nm laser (OBIS 405 nm LX 50 mW, Coherent).

The five laser lines were directed to the sample using a multi-band dichroic mirror (ZT405/488/561/647/752rpc-UF2, Chroma) on the excitation path. Laser intensities were controlled with an acousto-optic tunable filter (AOTF, 97-03309-01 Gooch & Housego), and laser on-off was controlled by mechanical shutters (LS3S2Z0, Vincent Associates). For cell line chromatin tracing experiments, we used a Lumencor CELESTA light engine for illumination, with the following laser wavelengths: 405-nm, 477-nm, 546-nm, 638-nm, and 749-nm. The lasers were directed to the sample using a corresponding penta-band dichroic mirror from Lumencor. Laser intensities and on-off were controlled by internal controls of the light engine. The 750/749-nm laser was used to excite and image DyLight 800-conjugated donkey anti-rabbit secondary antibody. The 647/638-nm laser was used to excite and image Alexa Fluor 647 (or Cy5) on readout probes and on the anti-rat secondary antibody. The 560/546-nm laser was used to excite and image tdTomato fluorescence and ATTO 565 (or Cy3) on readout probes. The 488/477-nm laser was used to excite and image GFP fluorescence and the yellow-green fiducial beads for drift correction. The 405-nm laser was used to excite and image the DAPI stain and the light-yellow fiducial beads. On the emission path, we had a multi-band emission filter (ZET405/488/561/647-656/752-nm Chroma for the tissue imaging setup or a corresponding penta-band emission filter for the cell line imaging setup) and a Hamamatsu Orca Flash 4.0 V3 camera. The pixel size of our system was 107.9 nm. To automatically scan and image multiple FOVs, we used a computer-controlled motorized x–y sample stage (SCAN IM 112×74, Marzhauser). For z-stepping and active auto-focusing, a piezo z positioner (Mad City Labs, Nano-F100S) was used.

### Arrayed RNAi screen of cell proliferation

Lentiviral infection conditions were optimized in 96-well plates for initial cell seeding number, lentiviral dosage, antibiotic concentration, and assay time. KP and *K-MADM-Trp53* LUAD cells were seeded in complete cell culture media at a density of 250 cells per well in a 96-well assay plate (Corning 3903), incubated for 24 hours, and infected with lentiviral supernatant of Sigma Mission^TM^ shRNAs targeting CPDs obtained from the Yale Cancer Center Functional Genomics Core. A total of 200 uL media was added into each well comprised of 50 uL lentiviral supernatant and 150 uL complete cell culture media with 10 ug/mL polybrene (EMD Millipore, TR-1003-G). Each shRNA hairpin was tested in triplicate. All lentiviral infections were performed in duplicate to rule out the influence of lentiviral dosage on cell growth: one replicate with 6 ug/mL puromycin selection and the other replicate with no puromycin selection. After 24 hours of lentiviral incubation, corresponding wells were treated with or without puromycin selection for 48 hours. The cells were then incubated with complete cell culture media for 4-5 days. Total viable cell count was determined with CellTiter-Glo Luminescent Cell Viability Assay (Promega, G7572) using a Promega luminescence plate reader. For data analysis, we deducted luminescence readout values of blank wells from those of test wells. We then normalized luminescence readout values of each target shRNA hairpin to those of shNTC. All shRNA hairpin sequences are provided in **Supplementary Table 6**.

### Rnf2 targeted degradation

Two million cells of the KP mouse primary adenocarcinoma cell line 31671 were nucleofected (Amaxa) with 2 μg homology-directed repair (HDR) template and 2 μM RNP (IDT Alt-R CRISPR-Cas9 sgRNA) for Rnf2-dTAG. The HDR template contained a 555-nt 5’ homology arm homologous to the 5’ end of the Rnf2 stop codon, a 30-nt linker and two HA tags, an FKBP^F36V^ and mScarlet insert (**Supplementary Table 6**, amplified from a gift plasmid from Dr. David Schatz at Yale University), and a 999-nt 3’ homology arm homologous to the 3’ end of the Rnf2 stop codon. After transfection, the cells were seeded into 6-well plates and cultured for 24-48 hours before FACS sorting. mScarlet+ cells were sorted into 96-well plates. Single colonies were manually picked, genotyped with PCR, and validated with sequencing. The spacer sequence for Rnf2 sgRNA is GACTTTATTATGCACCCACCA. The dTAG-13 ligand and negative control ligand were added to the cells at a final concentration of 500 nM. The cells were incubated at 37 °C with 5% CO_2_.

### Western Blot

Cells were trypsinized, harvested, and lysed with RIPA buffer (ThermoFisher, 89900) containing 1× protease inhibitors (ThermoFisher, 87786) at 4 °C for 30 min. Cell lysate supernatant was collected after centrifugation at 16,000 × g for 20 min at 4 °C. Supernatants were quantified using the BCA protein assay kit (Pierce, 23225). A total of 20 μg protein was denatured at 95 °C for 5 min and loaded on a 4-20% precast polyacrylamide gel (BioRad, 4568094) for gel electrophoresis. Proteins were transferred onto a PVDF membrane (ThermoFisher, IB24001) with an iBlot2 gel transfer device (Invitrogen, IB21001). The PVDF membrane was blocked with 5% (vol/vol) BSA in 1× TBST (AmericanBio, AB14330-01000), incubated with primary antibodies at 4 °C overnight, washed with 1× TBST for 5 min for three times, incubated with HRP-conjugated secondary antibodies at room temperature for 1 hour, and washed three times with 1× TBST for 5 min. Membranes were treated with SuperSignal West Pico Plus chemiluminescent substrate (ThermoScientific, 34577) and imaged with a ChemiDoc imaging system (BioRad). For fluorescence detection, proteins were transferred onto a nitrocellulose membrane (BioRad, 1620145) with the Trans-Blot Turbo transfer system (BioRad). Blots were washed once with 1x PBS (Boston BioProducts, BM-220X), blocked for an hour with Intercept Blocking Buffer (LiCOR, 927-7001), and incubated with primary antibodies overnight at 4°C. Blots were subsequently washed three times with 1x PBS-0.1%Tween20 for 10 min (Sigma-Aldrich, P1379-500ML), incubated with fluorescence secondary antibodies at room temperature for 45 minutes, wash three more times with 1x PBS-0.1%Tween20 followed by 1x PBS once prior to ChemiDoc imaging. The following antibodies were used: rabbit anti-Rnf2 (Cell Signaling Technologies, 5694S, 1:500), rabbit anti-Rnf2 (Proteintech, 16031-1-AP, 1:1000), mouse anti-Hsp90 (BD Biosciences, 610418, 1:10,000, HRP-conjugated goat anti-rabbit IgG (Abcam, ab6721, 1:3000), HRP-conjugated goat anti-mouse IgG (BioRad, STAR207P, 1:10,000), DyLight 800 4X PEG-conjugated goat anti-mouse IgG 800 (Cell Signaling Technologies, 5257S, 1:10,000), and DyLight 680-cojugated goat anti-rabbit IgG (Cell Signaling Technologies, 5366S, 1:10,000).

### Bulk RNA-sequencing of lung tumors

Large green tumors from *K-MADM-Trp53* mice were microdissected under a Nikon SMZ1270 fluorescence dissection stereo microscope and flash frozen. Tissue was pulverized using a BioPulverizer that was sprayed down with RNAse Away (Molecular BioProducts) and cooled with liquid nitrogen. RNA and genomic DNA were extracted using the AllPrep DNA/RNA Mini Kit (Qiagen). Library preparation for RNA-sequencing was performed by the Yale Center for Genome Analysis (YCGA), and libraries were sequenced on a NovaSeq S2 (Illumina) to obtain 100-bp paired-end reads. All reads that passed FASTQC quality metrics were mapped to the UCSC mm10 mouse genome and normalized gene count matrices were generated through STAR. Further analysis after trimming, alignment, and normalization were performed using DESeq2 on R^104^. Hierarchical clustering was done through the pheatmap package. Genes upregulated (log_2_ fold change > 2, FDR < 0.05) or downregulated (log_2_ fold change < -1, FDR < 0.05) in LUAD compared to AdenomaG were compared to the MSigDB Hallmarks gene set collection (https://www.gsea-msigdb.org/gsea/msigdb) to determine enrichment (hypergeometric test). Normalized expression counts and differential expression analyses are included in Supplementary Table 4.

### Whole exome sequencing analysis of copy number and single nucleotide variants

Tumor genomic DNA was obtained from flash frozen tumors using the AllPrep DNA/RNA Mini Kit, as described above. Paired normal DNA was obtained by extraction from FFPE slides of the same mouse lung by the YCGA. Whole exome sequencing (WES) was performed using the Mouse All Exon kit (Agilent) for target capture followed by next-generation sequencing by Psomagen. Mouse tumor samples were sequenced at 200× read coverage while healthy lung tissue was sequenced at 50×. For copy number analysis, sample reads were mapped to the GRCm38 reference genome with BWA-mem^105^, sorted based on coordinates with Picard SortSam tools (http://broadinstitute.github.io/picard/), and indexed with Samtools^106,107^. For single nucleotide variant (SNV) analysis, after initial quality control and trimming the raw sequences using fastp^108^, the trimmed sequence data were mapped to the mouse reference genome UCSC mm10 using BWA-MEM^105^. Duplicate reads were identified by employing the MarkDuplicates tool from the Genome Analysis Toolkit (GATK)/picard. Base Quality Score Recalibration (BQSR) was performed using BaseRecalibrator & ApplyBQSR with reference to the dbSNP database and data from the Sanger Mouse Genetics Programme (Sanger MGP). We created a panel of normals (PoN) containing germline and artifactual sites by running Mutect2 in tumor-only mode for each of 12 normal samples. We constructed a GenomicsDB datastore from the normal Mutect2 calls. The normal calls were combined to create the PoN with CreateSomaticPanelOfNormals. In this way, not only were the matched normal variants filtered out, but also any variants present in other normal mouse samples. We employed Mutect2 in GATK4 (v 4.4.0.0)^109^ to call somatic variants in a tumor/normal variant calling pipeline. The variant calling also utilized the PON file created using the normal calls. To address potential orientation biases in the raw data, we applied the LearnReadOrientationModel tool to learn the orientation bias model. Then, we filtered the unprocessed variants using FilterMutectCalls. We applied additional filtering criteria to reduce the false-positive rate of the variants identified. The positions in either tumor or matched normal samples with <10× coverage were removed from further analysis. At least three reads were required to support variants called in tumor samples, with no more than zero reads for the variant allele in the matched normal. A variant allele fraction of >5% was used to make mutation calls, which are listed in **Supplementary Table 7**.

### Single-nucleus RNA sequencing

#### Single-nucleus isolation and RNA sequencing library preparation

Nuclei from dissected lung tumors (two biologic replicates) were isolated by adapting a previously reported protocol^110^. Briefly, a stock solution of 2× salt-Tris buffer (ST buffer) composed of 292 mM NaCl (Thermo Fisher, BP358), 20 mM Trizma-HCl (Sigma, T2194-100ML), 2 mM CaCl2 (VWR, E506-100ML), and 42 mM MgCl2 (Alfa Aesar, J62411) in nuclease-free water (Invitrogen, 10977-15) was prepared fresh before isolation. 0.02% NP-40 Substitute based ST lysis buffer (NST lysis buffer) was generated using 1 mL of 2× ST buffer, 4 μL of 10% NP-40 Substitute (Sigma, 98379), 10 μL of BSA (NEB, B9000S), 20 μL of Superase-In RNase inhibitor (Invitrogen, AM2696), and 966 μL of nuclease free water (Invitrogen, 10977-015). Resuspension buffer was also freshly prepared using 880 μL of 1× Dulbecco’s PBS (Sigma, D8537-500ML), 100 μL of BSA, and 20 μL of Superase-In RNase inhibitor. 50 μL of NST lysis buffer was added to flash frozen tissue in a 1.5 mL microcentrifuge tube. The sample was continuously minced on ice with Noyes Spring scissors (Fine Science Tools, 15514-12) for four minutes to isolate nuclei. Additional NST lysis buffer was added to the sample for a final volume of 0.5 mL and passed through a 30 μm MACS SmartStrainer (Miltenyi Biotec, 130-098-458) into a 15 mL conical tube. The sample was then washed by adding 4 mL of 1x ST buffer through the strainer. Samples were spun down in a swinging bucket centrifuge for 5 min at 500 g at 4°C and resuspended in 50-100 μL of Resuspension buffer depending on the size of the pellet. The nuclei suspension was then passed through a 35***-***μm filter (Falcon, 352235). Nuclei were counted on a hemacytometer, and 10,000 nuclei were loaded onto a 10x Chromium chip for Chromium Single Cell 3’ Library (V3, PN-1000075) generation.

#### Single-nucleus RNA sequencing data analysis

Libraries were sequenced according to 10x Chromium manufacturer recommendations. The reads were aligned to the mm10-2020-A reference transcriptome to include introns using Cell Ranger count (v.7.1.0; 10x Genomics). To remove ambient RNA, raw matrices generated from Cell Ranger were inputted into Cellbender (Snapshot 11) using remove-background and run on the Terra platform with an FPR set to 0.01. Doublets were detected using Scrublet (v0.2.1) via doublet_detection (Snapshot 2) on Terra. Seurat_5.0.5 was used for downstream analyses. Cell barcodes with (1) 500-5000 genes; (2) 1000-10,000 transcript UMIs; and (3) less than 10% mitochondrial counts were included in the analysis. Data were normalized with a global scaling “LogNormalize” method and a scale factor of 10,000. We performed feature selection with the “vst” method and 2,000 features, scaled the data with all genes, performed principal component analysis (PCA) for dimensionality reduction, and clustered single nucleus gene expression with the Louvain algorithm. For tumor analyses, we first excluded fibroblasts and fibrocytes (*Ptprc, Cd163, S100a4, S100a8, S100a9, Cd90, Col1a1, Il6, Ccl3, Ccl4*), endothelial cells (*Pecam1, Cdh5, Tie2, Foxf1*), and immune cells (*Mrc1, Trac, Jchain, Ighg1*) and further identified adenoma cells (*Sftpc, Lyz2, Cxcl15, Hopx*), and LUAD cells (*Eif2s3y, Chsy3, Ldlrad4, Large1*) based on their marker gene expression patterns^40,110,111^.

### Image analysis

All image analysis was performed with MATLAB R2019b.

#### DAPI registration

GFP and tdTomato fluorescence images and SPC and CD45 co-immunofluorescence images were aligned to genome-wide chromatin tracing images using intensity-based image registration of the DAPI images. To process the DAPI images, we first took the average z-projection of each DAPI image stack and normalized it to its maximum intensity. We then reduced the background by normalizing the average projection image to the background calculated by the adaptthresh function. We then adjusted the threshold of the image so that the maximum and minimum intensities corresponded to the 3^rd^ and 1^st^ quartiles of the pixel intensities. We then applied an “opening-by-reconstruction” technique with a disk-shaped morphological structural element with a radius of 25 pixels to reduce the noise. Next, to align the processed DAPI images, we applied an intensity-based image registration algorithm. We used the imregtform function to estimate the geometric transformation for image alignment and the imregconfig function to generate the optimizer and metric configurations used by imregtform. We first optimized an initial transformation condition, and then used the optimized initial conditions to improve image alignment. For initial condition optimizations, we reduced the InitialRadius of the optimizer (generated by imregconfig) by a scale factor of 5 and set the MaximumIterations of the optimizer to 500. We then applied the optimizer and metric to the imregtform function with the “similarity” geometric transformation option to generate the initial geometric transformation object. We used the imregtform function to align the processed DAPI images with the “affine” geometric transformation option, the previously generated optimizer and metric, and the initial geometric transformation object. We finally generated a geometric transformation object to align DAPI images taken with fluorescent protein and co-immunofluorescence images to the DAPI images taken with the first hybridization round of genome-wide chromatin tracing.

#### GFP+ and tdTomato+ cell analysis

To identify cells with GFP or tdTomato fluorescence signals, we first generated maximum projections of GFP or tdTomato images along the z direction and aligned the images to the first-round readout hybridization images of genome-wide chromatin tracing. We then used an algorithm that can manually adjust the intensity threshold to determine GFP+ or tdTomato+ cells (Thresholding an image - File Exchange - MATLAB Central (mathworks.com)). We finally generated binary masks to distinguish GFP+ and tdTomato+ cells.

#### SPC+ and CD45+ cell extraction

To identify cells with SPC or CD45 immunofluorescence signals, we generated average projections of SPC or CD45 images along the z direction and normalized the images to the background calculated by the adaptthresh function. We then performed standard deviation filtering of the image with the stdfilt function, and filled holes with the imfill function, using a connectivity of 8 pixels. We then converted the SPC or CD45 images to binary images, used the regionprops function to identify SPC or CD45 patches, and excluded patches smaller than 150 pixels. The remaining patches were used to generated binary masks to distinguish SPC+ or CD45+ cells. Finally, to match the signals to nuclei, we generated binary masks for each cell nucleus using the DAPI images as described in the “Nucleus segmentation” section below and dilated the binary mask of each nucleus with a disk-shaped structural element of 10 pixels. We then multiplied each single-nucleus binary mask to the SPC or CD45 binary masks. Nuclei with more than 100 overlapping pixels were labeled as nuclei of SPC+ or CD45+ cells.

The genome-wide chromatin tracing image analysis pipeline consists of the following steps: color correction, drift correction, nucleus segmentation, foci fitting, decoding, and trace linking.

#### Color correction

The color shift between 647-nm and 560-nm laser channels was corrected by taking z-stack calibration images of Tetraspeck microspheres (0.1 μm, Invitrogen, T7279) attached to a coverslip surface. A polynomial spatial transformation structure in x and y was constructed with the cp2tform function and used for xy color shift correction. The color shift in z was corrected by calculating the mean z shift.

#### Drift correction

To correct for sample drifts between different rounds of hybridizations, we determined 3D positions (x, y, z) of fiducial beads with 3D Gaussian fitting for each hybridization round. We subtracted 3D positions (x, y, z) of the first hybridization round from each hybridization round to generate the drift correction profiles for all hybridization rounds.

#### Nucleus segmentation

Because the tissue sections largely consisted of a monolayer of cells, we segmented single cell nuclei in 2D based on DAPI staining patterns. We first applied drift corrections to the DAPI images and took their average projections along the z direction. We then normalized the DAPI average projection images to the background calculated by the adaptthresh function with a neighborhood size of 101 pixels. We further removed small “bright” objects using “opening-by-reconstruction” and small “dark” objects using “closing-by-reconstruction” techniques, both with a disk-shaped structuring element of 15 pixels. These processed DAPI images were further analyzed to extract foreground and background markers for the watershed algorithm. To obtain foreground markers for each single nucleus, we calculated the regional maxima with the imregionalmax function. To acquire background markers, we binarized the processed DAPI images and calculated their complement. We then used the imimposemin function to modify the processed DAPI images so that the regional minima occurred at foreground and background marker pixels. Finally, we applied the watershed function to the modified DAPI images for nucleus segmentation. We excluded small debris (<300 pixels) and under-segmented doublet nuclei (>9000 pixels) from our analyses.

#### Foci fitting

To determine the intensity threshold for DNA foci identification, we adapted a previously developed adaptive thresholding procedure so that the fitted foci number matched the expected DNA loci count^24^. The expected DNA loci count per nucleus per bit was approximately 20 based on the probe design. We then fitted 3D positions (x, y, z) of all DNA foci using a 3D radial center algorithm^112^ in each bit. We further applied color correction and drift correction to the fitted DNA foci in each bit, so that all fitted DNA foci were in the same 3D coordinate system as the first bit in the 560-nm laser channel. The signal intensity of each fitted DNA spot was normalized to the median signal intensities of all DNA foci in the corresponding image.

#### Decoding and trace linking

After we generated all fitted DNA foci in each bit in single nuclei, we adapted a previously reported expectation-maximization procedure for decoding^18^. First, we identified all valid spot pairs corresponding to a valid barcode whose two fitted DNA spots were within 500-nm spatial distance. For each spot pair, we calculated three quality metrics: (1) the distance between the 3D positions (x, y, z) of the spot pairs; (2) the difference between the signal intensities of the spot pairs; and (3) the average signal intensities of the spot pairs. We then calculated the percentages of spot pairs with worse qualities than a given spot pair (e.g. larger distance, larger difference, smaller intensities) and calculated the product of the three percentages as the quality score of the given spot pair. Next, for all spot pairs containing the same repetitive spot, we retained one spot pair with the highest quality score. Then, for each target genomic locus, we retained the top four spot pairs with the highest quality scores in each nucleus. After we obtained all processed spot pairs in all nuclei in each FOV, we linked the DNA loci positions into traces using a previously developed symmetric nearest neighbor approach^24^. We identified the 3D centroid positions of each linked chromosome territory and calculated a fourth quality metric: the distance between each spot pair to the nearest corresponding chromosome territory centroid. We then iteratively updated the quality scores, removed repetitive spot pairs, retained the top four spot pairs with the highest quality scores, and performed trace linking. For each single nucleus, if more than 99% (iteration rounds no more than 10) or 97% (iteration rounds more than 10) of spot pairs in the current iteration were the same as the previous iteration, the nucleus would be labeled as “decoded”. If more than 85% of nuclei in the FOV were labeled as “decoded”, the iteration would be terminated and spot pairs in the current iteration were stored as the finalized spot pairs. To link the finalized spot pairs (detected TADs) into chromatin traces, we first defined initial traces using the detected TADs in the first hybridization round. To grow the chromatin trace, we link the detected TADs to the traces if the detected TADs in the current hybridization round are the nearest neighbor to those in the previous hybridization round, and vice versa. After linking chromatin traces, we refit the missing TADs of each chromatin trace using the finalized spot pairs if they are within 6 pixels to the periphery of the chromosome territory. There are a small proportion (around 10%) of chromatin traces with overlapping TADs after the refitting procedure, so we further identified traces with >50% overlapping TADs and excluded the shorter ones. For each chromosome in a single cell, we then retained the longest two chromatin traces. Quality of the individual datasets was further confirmed by analyzing the detection efficiency of each target genomic region and trace length distribution, which was consistent across all datasets and all cancer states.

#### Fine-scale data analysis

For fine-scale chromatin tracing data analysis, the analytical procedure is the same as that in large-scale chromatin tracing mentioned above except for decoding and trace linking. We first fitted foci as described in “Foci fitting” above in 50 hybridization rounds in three color channels. The expected foci count per hybridization round was set as twice the number of target genomic loci in the specific round in the color channel. To link traces, we first identified the centroid position of the gene foci in the gene hybridization round, which approximated the centroid of the chromatin trace. Next, in each of the 40 genomic locus hybridization rounds, if a genomic locus was within 500 nm of the centroid, we included it into the chromatin trace. Chromatin traces shorter than 7 loci were removed. In each cell, we retained the longest two chromatin traces for each gene. To identify enhancer-promoter loops (E-P loops), we first calculated the mean spatial distance between each pair of loci. We excluded loci in hybridization rounds with poor signal-to-noise ratios. We then calculated the expected distances, as published previously^11,24^ using all cells of each state (power law fitting of mean spatial distance versus genomic distance). For each trace, we calculated the normalized distance (spatial distance/expected distance) between each pair of loci. We identified normalized distances between all loci to the promoter locus and performed one-sided Wilcoxon rank-sum tests and false discovery rate (FDR) multiple comparison correction to identify loci with lower normalized distances to the promoter than those of neighboring loci. If such locus contained a putative enhancer, we called it an E-P loop. We used E-P interactions in LUAD cells to identify CPD gene E-P interactions and those in AT2 plus adenoma cells to identify CTS E-P interactions. Putative enhancers were identified by the union of Ensembl predicted enhancers and H3K4me1 (ENCFF536DWZ) and DNaseI (ENCFF268DLZ) intersected ChIP-seq peaks in adult mouse lung.

### Data analysis and statistics

#### Heterogeneity

To compare chromatin conformational heterogeneity, we defined a heterogeneity score as the coefficient of variation (COV) of inter-loci distances between a pair of TADs on a chromosome of a cell state (variation among different copies of the chromosome). Wilcoxon signed-rank test was used to compare all heterogeneity scores of a chromosome between cell states.

#### Decompaction

To compare levels of chromatin compaction, we defined a decompaction score as the mean inter-loci distance between a pair of TADs on a chromosome of a cell state. Wilcoxon signed-rank test was used to compare all decompaction scores of a chromosome between cell states.

#### Demixing

To compare levels of chromatin intermixing or demixing, we defined a demixing score as the standard deviation of all normalized mean inter-loci distances on a chromosome of a cell state. The normalized mean inter-loci distance between a pair of TADs was calculated by normalizing the mean inter-loci distance between the pair of TADs to the average value of all mean inter-loci distances on the chromosome. Levene’s test of normalized mean inter-loci distances of a chromosome was used for statistical comparisons between cell states for potential differences in chromatin intermixing/demixing.

#### A and B compartment polarization analysis

To identify A and B compartments, we adapted a previously developed algorithm to determine A and B compartment scores^11,24^. In brief, we first generated a mean inter-loci spatial distance matrix for each chromosome, where each pixel corresponded to the mean spatial distance between a pair of TADs. We then normalized the mean spatial distances to the corresponding expected spatial distances calculated by power-law fitting of spatial versus genomic distances. Next, we calculated the Pearson correlation coefficient between each pair of columns of the normalized matrix. Finally, we applied a principal component analysis of the Pearson correlation matrix and used the coefficients of the first principal component as the compartment scores. We calculated the correlations between compartment scores and an averaged profile of H3K4me1, H3K4me3, DNase I hypersensitivity site, and gene densities^113,114^, and flipped the signs of the compartment scores if necessary, so that the correlations were positive. Under this assignment, A compartment TADs had positive scores and B compartment TADs had negative scores. We then used a previously described polarization index metric to quantify the polarized arrangement of A and B compartments ^24^. We downloaded the called H3K4me1 and H3K4me3 ChIP-seq peaks and DNA DNase I hypersensitivity sites from the ENCODE portal with the following identifiers: ENCFF536DWZ, ENCFF508WEP, ENCFF268DLZ. The gene density profile was downloaded from the UCSC table browser.

#### Radial score analysis

To calculate the radial score of each TAD in each single cell, we first measured the mean spatial distance between each TAD to the centroid of all target TADs and normalized the distance to the average spatial distances from all TADs to the centroid. Wilcoxon signed-rank test was used to compare all radial scores of TADs on a chromosome between cell states.

#### The ”Trace2State” pipeline for single-cell chromatin conformation-based cell state visualization and classification

To visualize in low dimension potential clustering of single-cell 3D genome conformations, we constructed an input matrix where each row corresponded to a single cell and each column represented the single-cell A/B compartment (scA/B) score of each TAD^34^. Only cells with at least 10 traces were analyzed. The scA/B score of each TAD was calculated as the mean A/B compartment score of all its spatially adjacent TADs within a 1200-nm 3D radius neighborhood. Specifically, we first calculated the population-average A/B compartment scores of each TAD as described in “*A and B compartment polarization analysis*” above. For each observed TAD locus, we then identified all other TADs within a 1200-nm 3D radius neighborhood and calculated their mean A/B compartment score as the scA/B score of the central TAD locus. Missing values in the matrix were replaced with 0’s. The matrix was used as input in a principal component analysis, and the first 50 principal components were used for downstream visualizations. We scaled the 50 principle components with the PCA().fit-transform function in python, and visualized the single cell data with t-distributed stochastic neighbor embedding (t-SNE), UMAP and PacMAP^36–38^. To classify different cell states with supervised machine learning, we supplied the Classification Learner App of MATLAB with the same input scA/B score matrix as described above. To evaluate the supervised machine learning model, we applied five-fold cross-validation which prevents overfitting. Specifically, data were partitioned into five randomly chosen subsets of roughly equal sizes, with four assigned as training subsets and one as the test subset. The model was trained repeatedly five times such that each subset was used exactly once as the test subset. The overall prediction accuracy was calculated as the average prediction accuracy of the test subsets in the five trainings. The Median Gaussian SVM model we used in MATLAB Classification Learner included regularization (box constraint level = 1). All machine learning models were trained, and the model with the highest prediction accuracy (medium gaussian support vector machine) was retained to plot the confusion matrix and ROC curves.

#### The “Trace2Biomarker” pipeline for CPD and CTS gene identification

To identify CPD and CTS genes, we first identified marker TADs with significantly changed scA/B scores during LUAD progression. Specifically, we extracted cells with more than 325 TADs. We next performed rank-normalization using the tiedrank function in MATLAB of scA/B scores in each cell. We performed Wilcoxon rank-sum tests of rank-normalized scA/B scores of each TAD comparing AdenomaG and LUAD cells. TADs with *p* < 0.1 were defined as marker TADs. We then defined CPD genes as genes located in marker TADs with increased scA/B scores and with significantly higher expression (mean expression count > 10, fold change > 3, FDR < 0.05) in LUAD compared to AdenomaG cells. CTS genes were defined as genes located in marker TADs with decreased scA/B scores and with significantly lower expression (mean expression count > 10, fold change < 0.5, FDR < 0.05) in LUAD compared to AdenomaG cells. For control groups, genes with significantly increased expression in regions with unchanged scA/B scores are defined as those in TADs with p >= 0.1 and increased expression (mean expression count > 10, fold change > 3, FDR < 0.05) in LUAD compared to AdenomaG cells. Genes with significantly decreased expression in regions with unchanged scA/B scores are defined as those in TADs with p >= 0.1 and decreased expression (mean expression count > 10, fold change < 0.5, FDR < 0.05) in LUAD compared to AdenomaG cells. Genes with increased expression only are defined as those with increased expression (mean expression count > 10, fold change > 3, FDR < 0.05) in LUAD compared to AdenomaG cells irrespective of scA/B score changes. Similarly, genes with decreased expression only are defined as those with decreased expression (mean expression count > 10, fold change < 0.5, FDR < 0.05) in LUAD compared to AdenomaG cells. Equal numbers of genes (the top 21 upregulated and top 19 downregulated) were used for each of the above gene lists interrogated in TCGA patient survival analyses described below.

#### Quantification of single-nucleus and single-cell gene expression homogeneity

For expression homogeneity calculation, we used a similar algorithm as described previously^40^. We first randomly subsampled 100 cells and calculated the Pearson correlation coefficient of gene expression between each cell pairs. We repeated the process 100 times, each time using the mean Pearson correlation coefficient as the gene expression homogeneity score.

#### Identification of AT2 cells spatially close to and far from immune cells

To distinguish AT2/cancer cells close to and far from immune cells, we extracted binary masks of nuclei of SPC+ cells (or GFP+CD45− cancer cells in LUAD tumors) and of CD45+ cells and dilated the SPC+/cancer cell binary mask by 50 pixels. The SPC+/cancer cells with overlapping pixels with the CD45+ mask were identified as AT2/cancer cells spatially close to immune cells, whereas SPC+/cancer cells with no overlapping pixels with the CD45+ mask were identified as AT2/cancer cells distant from immune cells.

#### Probability for randomly selected genes to inhibit cell growth

To quantify the percentage of randomly selected genes that can affect cell growth, we downloaded the gene_dependency.csv table from https://figshare.com/articles/dataset/DEMETER_2_Combined_RNAi/9170975, which contained the probabilities that knocking down one gene has a cell growth inhibition or death effect. Cancer cell line genetic dependencies were estimated using the DEMETER2 model applied to a combination of three large-scale Cancer Dependency Map RNAi screening datasets (the Broad Institute Project Achilles, Novartis Project DRIVE, and the Marcotte et al. breast cell line dataset). We identified all mouse LUAD cell lines from the table with available RNAi data and calculated the mean probability of all genes in mouse LUAD cells. 6.4% of randomly selected genes are expected to affect cell growth.

#### TCGA patient survival analysis

Clinical data for TCGA LUAD patient survival and RNA-seq data were obtained from the GDAC website from the Broad Institute (https://gdac.broadinstitute.org/). The identified CPD/CTS genes of LUAD were converted to human orthologs. The RNA-seq expression matrix and the gene list were applied as input to score the individual expression files using GSVA R package with ssGSEA scoring method^115,116^. TCGA LUAD patients were grouped based on high/low ssGSEA scores (most correlated/least correlated) using the top/bottom quintiles. Kaplan-Meier survival analysis was carried out using survfit R function with log-rank significance test.

#### The “Trace2Regulator” pipeline for 3D genome regulator identification

To identify chromatin regulators that bind to genes in marker loci, we first identified genes with increased expression (FDR < 0.1, fold change > 1) in TADs with increased scA/B scores (p < 0.1) from adenoma green to LUAD cells. We then inputted the gene list into the BART algorithm^51,52^ to generate a list of chromatin regulators predicted to bind to the input genes, using the default parameter settings. We further knocked down candidate chromatin regulators with shRNAs in the KP LUAD cell line and performed DNA MERFISH. We further compared 3D genome alterations upon the candidate chromatin regulator knockdown with those during the adenomaG to LUAD progression. Candidate regulators showing concordant 3D genome alterations were identified as 3D genome regulators.

### CUT&RUN experiments and analysis

#### CUT&RUN protocol

CUT&RUN was performed using the Epicypher CUTANA^TM^ ChIC/CUT&RUN Kit (14-1048) according to the manufacturer’s specifications with the following conditions and modifications: For binding to the Concanavalin beads, 500,000 cells per sample were prepared, counted twice by hemacytometer, and averaged. Nuclei were prepared according to the Epicypher Cut and Run Manual Appendix. Nuclei were incubated with activated beads. 0.1 *E. coli* spike in DNA ng was added to each sample. The following antibodies (0.5 ug of antibody per reaction) were used:

IgG Control antibody: CUTANA^TM^ Kit Rabbit IgG CUT&RUN Negative Control Antibody

RNF2 antibody: CST RING1B (D22F2) XP® Rabbit mAb #5694

H3K4me3 antibody: Epicypher Rabbit Polyclonal H3K4me3 13-0041

H3K27me3 antibody: CST Tri-Methyl-Histone H3 (Lys27) (C36B11) Rabbit mAb #9733

H2AK119ub antibody: CST Ubiquityl-Histone H2A (Lys119) (D27C4) Rabbit mAb #8240

BMI-1 antibody: Active Motif BMI-1 antibody (mAb) 39993

RNA Pol II p-ser5: Abcam Anti-RNA polymerase II phosphor-S5 EPR19015

#### Library preparation

Libraries were prepared using the Epicypher CUTANA^TM^ CUT&RUN Library Prep Kit (14-1001) according to the manufacturer’s specifications with the following modification: SPRI-select beads were used after library preparation to perform 200-700 bp size selection to enrich for DNA fragments from CUT&RUN and remove any adapter dimers or high molecular weight DNA. Library quality was analyzed using Agilent Tapestation D1000 High Sensitivity Tapes (#5067-5584). Sequencing was performed with 10 million reads ordered per sample, 150 bp paired-end reads, on an Illumina NovaSeq 6000.

#### CUT&RUN data analysis

CUT&RUN reads were analyzed for quality using FastQC (https://www.bioinformatics.babraham.ac.uk/projects/fastqc/) and trimmed using Trimmomatic. Reads were then aligned to mm9 and K-12 *E. coli* genome U00096.3 by Bowtie2 version 2.3.4. Picard version 2.27.4 was used to down-sample the reads so that the read depths of *E. coli* sequences from different samples matched each other. SAMTools version 1.11 was then used to convert to BAM format, index, isolate uniquely mapped paired reads, and remove duplicates.

MACS2 version 2.2.7.1 was used to call sample narrow peaks, using IgG as an input. Read counts across genomic intervals and peak visualization were performed using deepTools version 3.3^117^, and .bw files were visualized using the Integrative Genomics Viewer.

## Data availability

All raw imaging data are available upon request. All sequencing data will be available at the Gene Expression Omnibus (GEO) upon publication. Analyzed imaging and sequencing data are available at https://campuspress.yale.edu/wanglab/Cancer3DGenome/.

## Code availability

Open-source code for imaging data collection is available at https://github.com/ZhuangLab/storm-control. MATLAB code for raw image analysis and downstream data analysis are available at https://campuspress.yale.edu/wanglab/Cancer3DGenome/.

## Acknowledgements

We thank Drs. Andrew Xiao, Megan King, Bogdan Bintu, Pu Zheng, Robert Homer, Katerina Politi, Roy Herbst, and members of the Wang and Muzumdar labs for helpful discussions, Dr. Mark Lemmon for critical reading of the manuscript, the Yale Cancer Center (YCC) Functional Genomics Core for assistance with RNAi screening assays, Yale Pathology Tissue Services (YPTS) for histology support, Drs. Jeffrey Ishizuka and David Braun for generously offering lab space for nuclei isolation for snRNA-seq, the Yale Center for Genome Analysis (YCGA) for RNA and snRNA library preparation and sequencing of wild-type lung and mouse tumors, Dr. Arjun Bhutkar and Sarah Blatt for sharing the ssGSEA computational pipeline for TCGA survival analyses, the ENCODE Consortium and the labs of Drs. Bing Ren and John Stamatoyannopoulos for generating the H3K4me1 and H3K4me3 ChIP-seq and DNase-seq datasets in adult mouse lung, Dr. Nik Joshi for providing the KP cell line, Drs. Robert Weinberg, Didier Trono, and David Schatz for constructs, and Drs. Tyler Jacks and Liqun Luo for mouse strains. S.S.A. and C.F.R. were supported by postdoctoral fellowships through the Yale Cancer Biology Training Program (T32CA193200). C.F.R. is supported by an NIH Research Supplement to Promote Diversity in Health-Related Research (R01CA276108-01A1S1). T.B.J. was in part supported by 5T32GM007205. J.S.D.R. was supported by a NIH Predoctoral Training Grant (2T32GM007499). J.P.T. recognizes support from NIH (R01LM013385). M.D.M. acknowledges support from a NIH Director’s New Innovator Award (DP2CA248136), AACR-Genentech NextGen Grant for Transformative Cancer Research (19-20-18-MUZU), Yale SPORE in Lung Cancer Career Enhancement Program award (P50CA196530), and in part, the Yale Comprehensive Cancer Center Support Grant (P30CA016359). S.W. was partly supported by the NIH (UH3CA268202, U01CA260701, R01HG011245, DP2GM137414, R01HG012969), Pershing Square Sohn Cancer Research Alliance, American Federation for Aging Research and Hevolution Foundation. This work was largely supported by an NCI IMAT program award (R33CA251037) and R01CA292936 to M.D.M. and S.W.

## Author contributions

M.D.M. and S.W. conceived of and supervised the study. M.L., M.D.M., and S.W. designed the study with help from S.J. and S.S.A. M.L., S.J., S.S.A., T.B.J., and J.S.D.R. performed experiments. M.L., S.J., S.S.A., T.B.J., T.Y. C.F.R., G.B, M.R. analyzed data with help from J.P.T., M.D.M., and S.W. M.L., S.J., S.S.A., T.B.J., M.D.M. and S.W. wrote the manuscript with input from all authors.

## Competing interests

S.W., M.L., M.D.M., S.J., and S.S.A. are inventors on a patent applied for by Yale University related to this work. M.D.M. received research funding from a Genentech supported AACR grant and an honorarium from Nested Therapeutics. The remaining authors declare no competing interests.

## Supplementary information

**Supplementary Table 1. Template oligonucleotide probe libraries for genome-wide and fine-scale chromatin tracing**

**Supplementary Table 2. Codebook for all target genomic loci for genome-wide chromatin tracing.** The first column indicates which chromosome each target TAD is located in. The second and third column indicate the start coordinate and end coordinate of the target TAD. The fourth column indicates the HW2 binary barcode for each TAD. Each barcode consists of 100 bits, with 2 bits assigned as ‘1’ and all the other 98 bits as ‘0’.

**Supplementary Table 3. Oligo sequences for adapters, common readout probes, blocking oligos and primers.** The file contains four spreadsheets: ‘Large-scale adapters’, ‘Fine-scale adapters’, ‘Common readout oligos’, ‘Blocking oligos’ and ‘Primers’. ‘Large-scale adapters’ contains the sequences of all 100 oligos that can bind to the primary probes and correspond to the 100 bits in the codebook. ‘Fine-scale adapters’ contains the sequences of loci-specific probes and gene-specific probes. ‘Common readout oligos’ contains the sequences of dye-conjugated readout probes. ‘Blocking oligos’ contains the dye-free oligos that have the same sequences as the common readout oligos to block any unbound sites of the previous adapters. ‘Primers’ contains the forward and reverse primer sequences for the template probe library amplification and primary probe synthesis.

**Supplementary Table 4. RNA-seq analysis of lung tumors from *K-MADM-Trp53* mice.** Differential expression analysis (DESeq2) of large green tumors derived from *K-MADM-Trp53* mice and classified as AdenomaG and LUAD based on marker gene expression. For each gene, log2 normalized expression counts (DESeq2) for each sample, average expression counts across all samples, log2 fold change (LUAD vs. AdenomaG), p-value, and FDR (padj) are shown.

**Supplementary Table 5. List of candidate progression driver, candidate tumor suppressor and candidate tumor initiation genes.** Lists of candidate progression drivers, candidate tumor suppressors, and candidate tumor initiation genes. References (PubMed IDs (PMID)) are listed for genes with prior literature evidence for a functional role in lung cancer pathogenesis.

**Supplementary Table 6. shRNA sequences for lentiviral transduced stable cell lines to knock down candidate progression driver genes and sequence of the FKBPF36V and mScarlet insert.** The table contains two spreadsheets. Spreadsheet one contains sequences for all the shRNAs of the candidate genes, positive controls, and negative controls. The first column contains the gene names. The second column contains the shRNA sequences. The third column contains the hairpin numbers. Spreadsheet two contains sequences for the FKBPF36V and mScarlet insert used for Rnf2 targeted degradation.

**Supplementary Table 7. Whole exome sequencing analysis of lung tumors from *K-MADM-Trp53* mice.** Mutation calls of variants (variant allele fraction > 5%) for adenoma and LUAD tumors (n = 6 tumors per group). Table includes mouse number (sample), variant chromosome, genomic position, reference base (REF), variant base (ALT), variant allele fraction (Tumor_AF), number of reference read (AD[0]) and variant reads (AD[1]) in tumor and normal, gene, and protein-altering effect (missense, splice variant, stop_gained).

## Extended Data Figures

**Extended Data Fig. 1.**
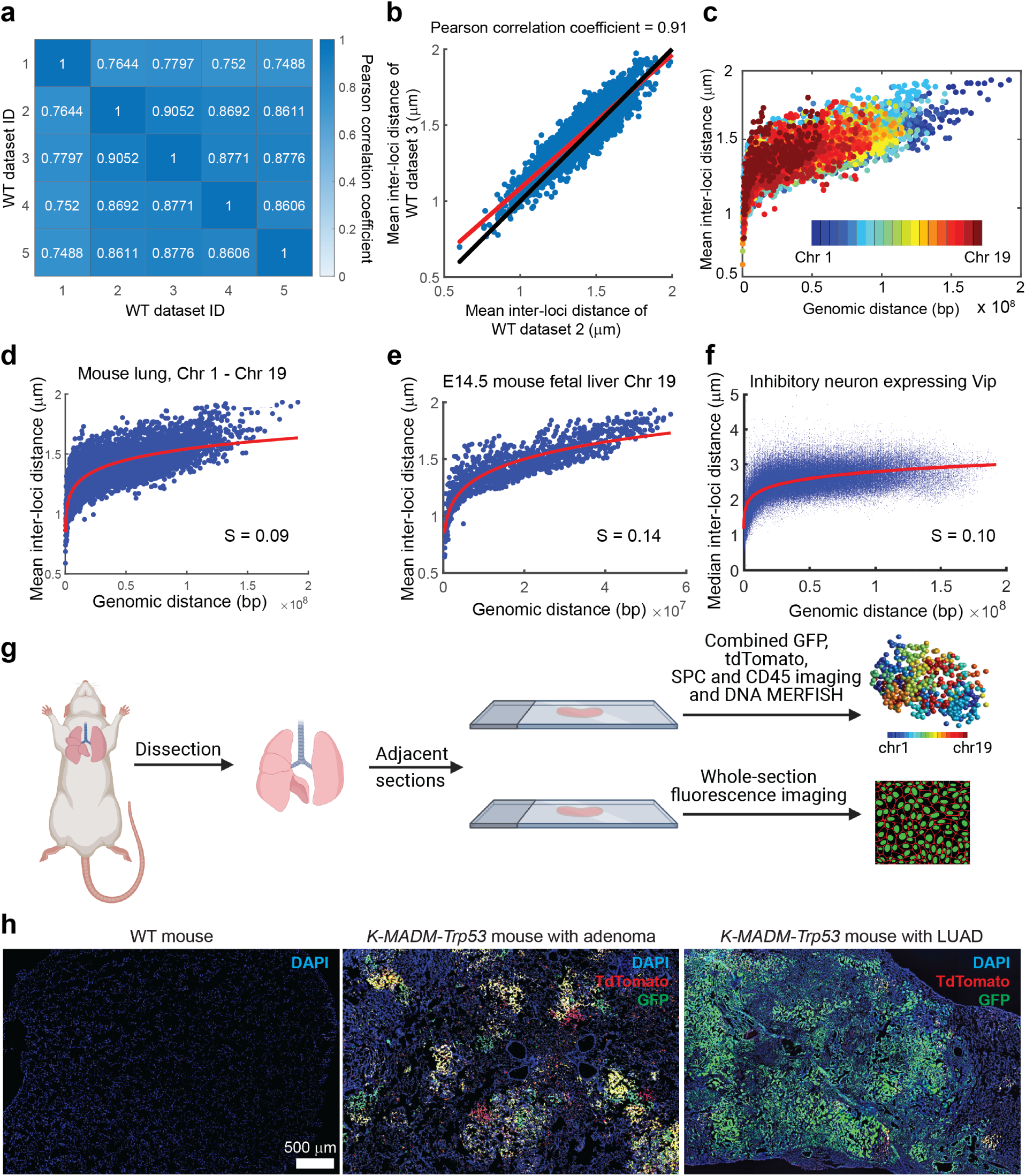
Genome-scale chromatin tracing visualizes 3D genome organization *in vivo*. **a,** Pearson correlation coefficients of mean inter-loci distances between WT datasets. **b,** Mean inter-loci distances of WT dataset 3 versus mean inter-loci distances of WT dataset 2. The red line is a fitted linear regression line. The black line is the y = x line. **c,** Mean inter-loci spatial distance versus genomic distance for all pairs of genomic loci on each autosome in AT2 cells. Different pseudo-colors represent different autosomes. n = 6,039 intra-chromosomal inter-loci pairs in **b-d**. n = 4,806 WT AT2 cells in **c-d**. **d,** Power-law scaling of all 19 mouse autosomes (Chr 1-19) in WT mouse lung. **e,** Power-law scaling of Chr 19 in E14.5 mouse fetal liver. Data were re-analyzed from Liu et al. Nat. Commun. (2020)^24^. **f,** Power-law scaling of all 20 mouse chromosomes (Chr 1-19, Chr X) in the mouse brain inhibitory neurons expressing Vip. Data were re-analyzed from Takei et al. Science. (2021)^23^. **g,** Schematic illustration of the experimental procedure. The schematic is created with BioRender.com. **h,** Whole-section fluorescence images of wild-type (WT) mouse lung and *K-MADM-Trp53* mouse lungs containing adenomas or LUAD. For panels **d-f**, lines are fitted power-law functions, and S is the scaling factor.

**Extended Data Fig. 2.**
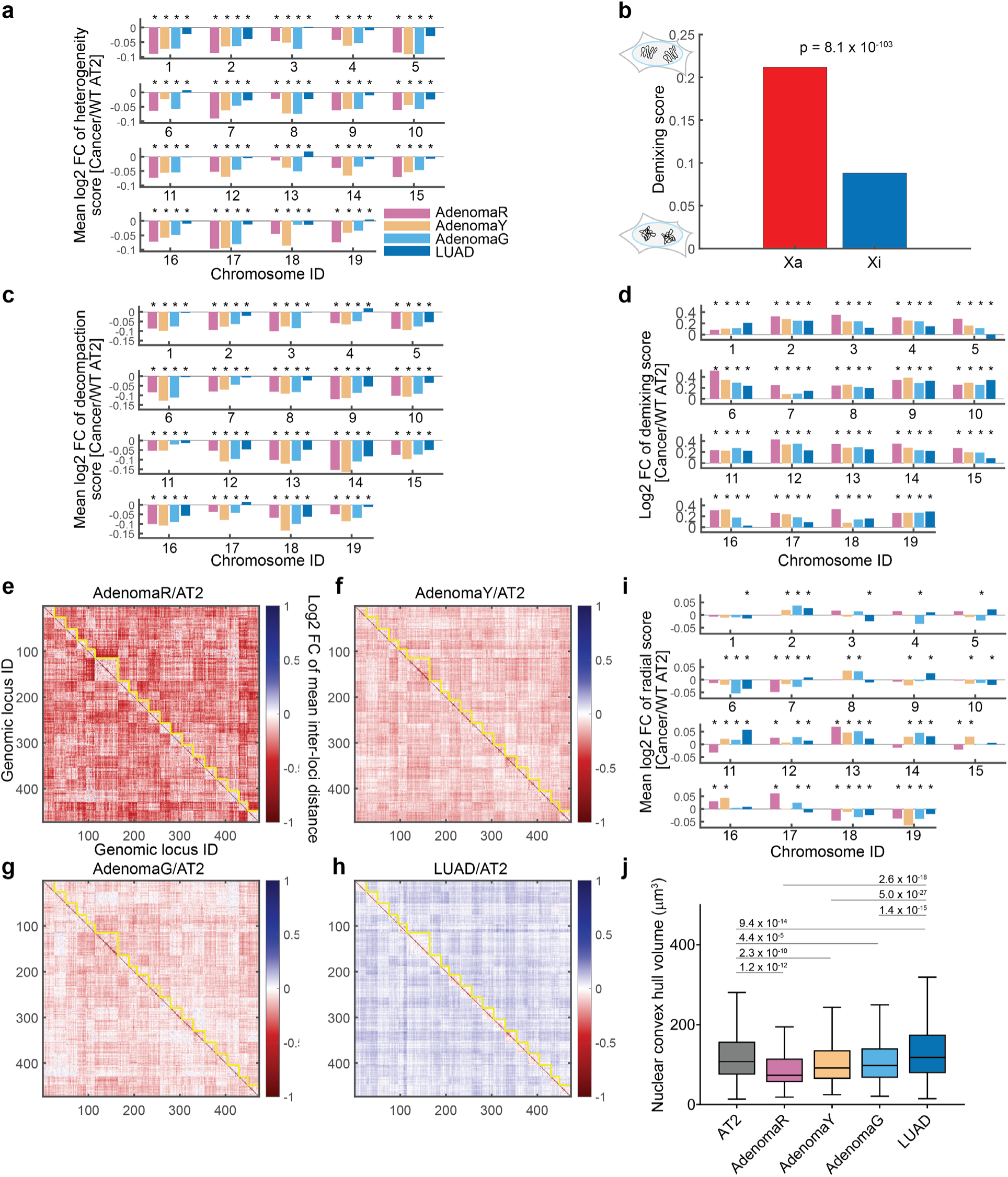
Changes of cancer state-specific 3D genome features comparing each cancer state to the AT2 state. **a,** Mean log2 fold change of heterogeneity scores of each chromosome, comparing each cancer state to WT AT2 cell state. * indicates FDR < 0.05, two-sided Wilcoxon signed-rank test. **b,** Demixing scores (standard deviation of normalized mean inter-loci distances) of the active X chromosomes (Xa, n = 95) and inactive X chromosomes (Xi, n = 95) in human IMR90 cells show a reduction (increased intermixing) in Xi, as previously described using other analyses^11^. p value from two-sided Levene’s test is shown. **c,** Mean log2 fold change of decompaction scores of each chromosome, comparing each cancer state to WT AT2 cell state. * indicates FDR < 0.05, two-sided Wilcoxon signed-rank test. **d,** Log2 fold change of the demixing score of each chromosome, comparing each cancer state to WT AT2 cell state. * indicates FDR < 0.05, two-sided Levene’s test. **e-h,** Log2 fold changes of mean inter-loci distances for adenoma red (AdenomaR) (**e**), adenoma yellow (AdenomaY) (**f**), adenoma green (AdenomaG) (**g**), and LUAD (**h**) relative to AT2 cells. Yellow lines highlight the boundaries of chromosomes. **i,** Mean log2 fold change of radial scores of each chromosome, comparing each cancer state to the AT2 cell state. * indicates FDR < 0.05, two-sided Wilcoxon signed-rank test. **j,** Nuclear convex hull volume of WT AT2, AdenomaR, AdenomaY, AdenomaG, and LUAD cells. p values of two-sided Wilcoxon rank-sum tests are shown. The horizontal lines of each box from top to bottom represent the 75th percentile, median, and 25th percentile. Whiskers extend to the non-outlier maximum and non-outlier minimum. Outliers are defined as values at least 1.5 times interquartile range away from the top or bottom of the box. Cell numbers in (**a, c-j**) are the same as in Figure 2.

**Extended Data Fig. 3.**
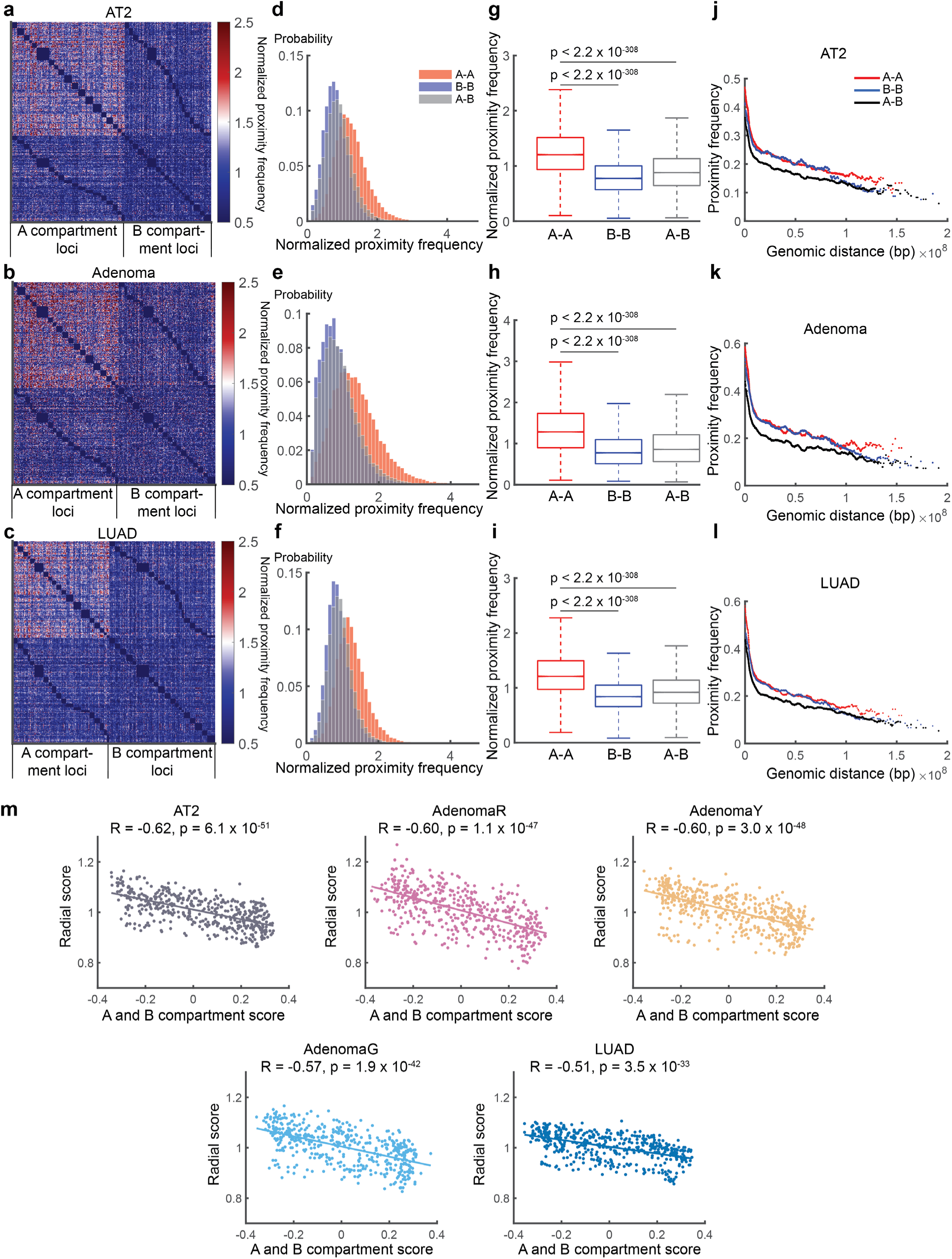
Cancer state-independent 3D genome features during lung cancer progression. **a-c,** Normalized trans-chromosomal proximity frequency between genomic loci in AT2 (**a**), adenoma (**b**), and LUAD (**c**) cells. The proximity frequency between each pair of trans-chromosomal genomic regions were normalized to the mean proximity frequency of all loci pairs of the two corresponding chromosomes. A cutoff distance of 800-nm was used for defining proximity. The genomic regions were re-ordered so that A compartment loci were grouped separately from B compartment loci. **d-f,** Distribution of normalized trans-chromosomal proximity frequencies of pairs of A loci (A-A), pairs of B loci (B-B) and pairs of A and B loci (A-B) in AT2 (**d**), adenoma (**e**) and LUAD (**f**) cells. **g-i,** Normalized trans-chromosomal proximity frequencies of A-A, B-B and A-B loci pairs in AT2 (**g**), adenoma (**h**) and LUAD (**i**) cells. The horizontal lines of each box from top to bottom represent the 75th percentile, the median and the 25th percentile. Whiskers extend to the non-outlier maximum and non-outlier minimum. Outliers are defined as values at least 1.5 times interquartile range away from the top or bottom of the box. p values from two-sided Wilcoxon rank-sum test are shown. **j-l,** The proximity frequency between each pair of cis-chromosomal genomic regions as a function of their genomic distances in AT2 (**j**), adenoma (**k**), and LUAD (**l**) cells. **m,** Radial scores versus A-B compartment scores of genomic loci in the five cell states. The lines are fitted linear regression lines. Correlation coefficients (R) and p values are shown. Cell numbers for all panels are the same as in Figure 2.

**Extended Data Fig. 4.**
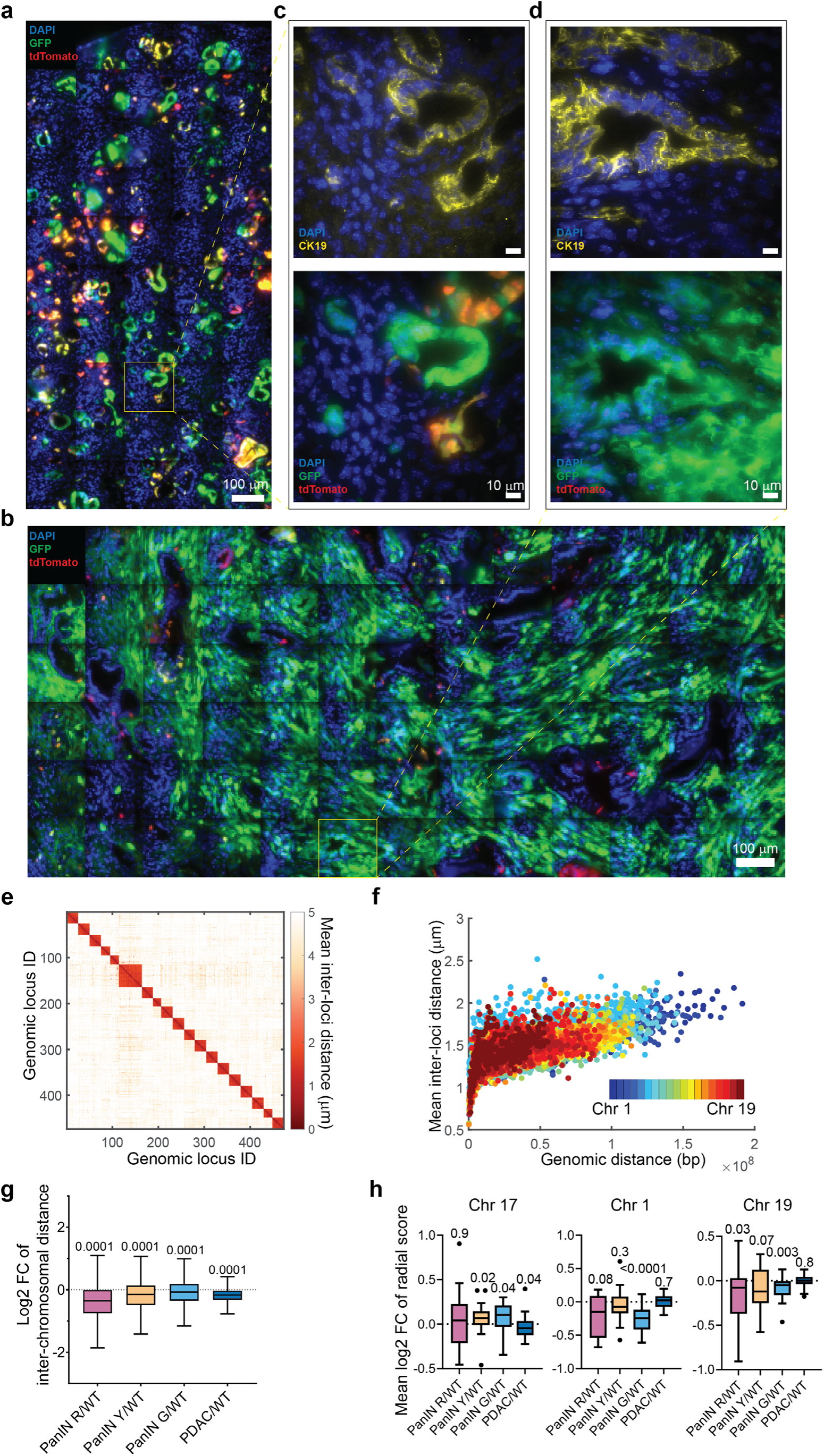
Genome-wide chromatin tracing of pancreatic adenocarcinoma progression. **a-b,** Whole-section fluorescence images of a K-MADM-Trp53 pancreas with PanINs (**a**) and PDAC (**b**). **c-d,** (Upper panels) Immunofluorescence staining of the duct cell marker CK19 and DAPI staining in a field of view in **a-b**. (Lower panels) Fluorescent protein imaging of GFP and tdTomato and DAPI staining in a field of view in **a-b**. The images are maximum-intensity z-projections from 10-μm z-stacks. **e,** Matrix of mean inter-loci distances between all genomic loci in normal duct cells. n = 1,529 cells. **f,** Mean inter-loci spatial distance versus genomic distance for all pairs of genomic loci on each autosome in duct cells. Different pseudo-colors represent different autosomes. n = 6,039 intra-chromosomal inter-loci pairs. n = 1,529 cells. **g,** Log2 fold change of mean inter-chromosomal distances, comparing each cancer state to normal duct cells. n = 1529, 123, 189, 361, 475 for normal duct, PanIN R, PanIN Y, PanIN G, and PDAC cells. Cell numbers of each cell state are identical in **g** and **h**. **h,** Distribution of the mean log2 fold change of radial scores of Chr17, Chr1, and Chr19, comparing each cancer state to normal duct cells.

**Extended Data Fig. 5.**
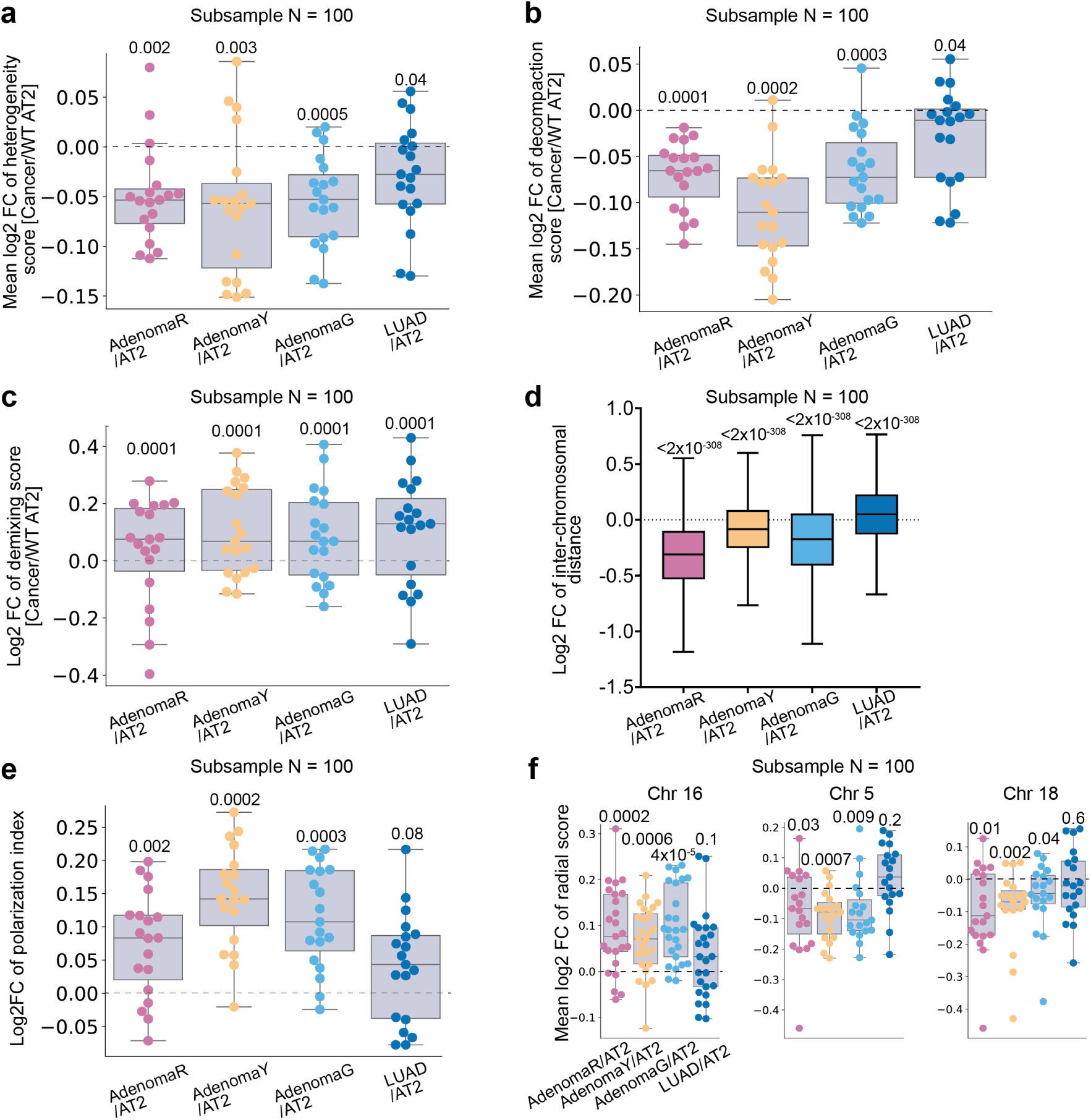
Subsampling analysis of chromatin folding changes during lung cancer progression. **a-c,** Distributions of the (mean) log2 fold change of heterogeneity (COV of inter-loci distance), decompaction (mean inter-loci distance), and demixing scores of each autosome (n = 19) in a randomly subsampled population of 100 cells per cell state, comparing each cancer state to AT2 state. **d,** Distribution of the log2 fold change of inter-chromosomal distances in a randomly subsampled population of 100 cells per cell state, comparing each cancer state to AT2 state. **e,** Distribution of the log2 fold change of polarization indices of A and B compartments of each autosome (n = 19) in a randomly subsampled population of 100 cells per cell state, comparing each cancer state to AT2 state. **f,** Distribution of the mean log2 fold change of radial scores of Chr 16, Chr 5, and Chr18 in a randomly subsampled population of 100 cells per cell state, comparing each cancer state to AT2 state. p values of two-sided Wilcoxon signed-rank test (**a-f**) are displayed. In **a-f**, the horizontal lines of each box from top to bottom represent the 75th percentile, median, and 25th percentile. Whiskers extend to the non-outlier maximum and non-outlier minimum. Outliers are defined as values at least 1.5 times interquartile range away from the top or bottom of the box.

**Extended Data Fig. 6.**
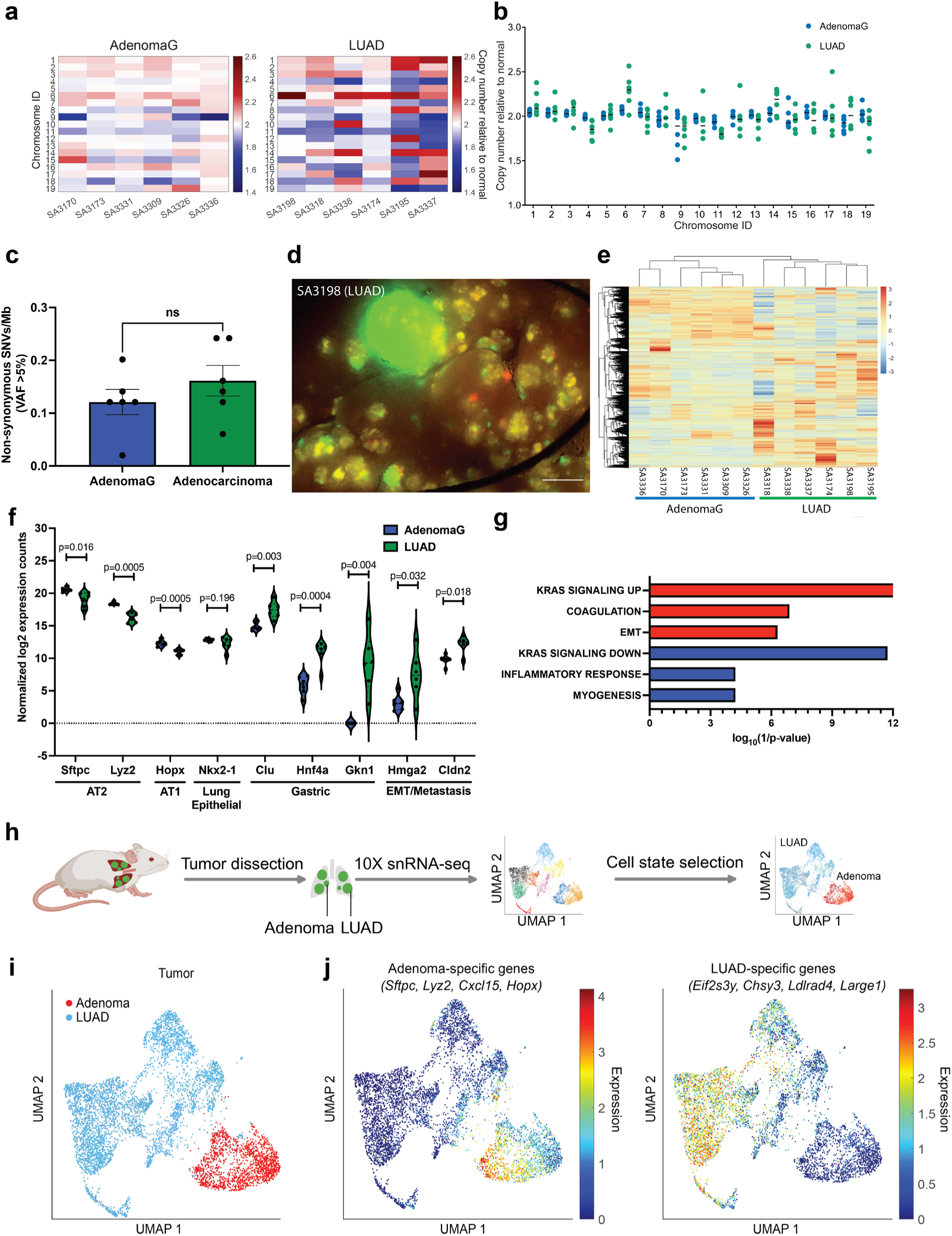
Exome and RNA sequencing analyses of lung tumors from *K-MADM-Trp53* mice. **a,** Heatmap of mean copy number variants (CNVs) of each autosome across AdenomaG and LUAD samples (n = 6 tumors per group). Classification of AdenomaG and LUAD is based on parallel gene expression analysis by RNA-seq on the same tumors (**e**). **b,** Distribution of copy numbers of each autosome in AdenomaG and LUAD tumors relative to paired normal. Each dot represents a single tumor. Black lines represent the median values. **c,** Frequency of non-synonymous single nucleotide variants (SNVs) per megabase (Mb) with variant allele fraction (VAF) > 5% in AdenomaG and LUAD tumors. **d,** Representative large GFP+ (green) tumor dissected under fluorescence microscopy for whole exome and RNA sequencing analyses. Scale bar = 2.5 mm. **e-f,** Unsupervised hierarchical clustering of all expressed genes (**e**) segregates dissected green tumors into two clusters (n = 6 tumors per cluster) defining AdenomaG and LUAD cells based on the expression of previously described markers of histologic progression^40^ (**f**), including loss of AT2/AT1 genes (Stfpc, Lyz2, and Hopx) and acquisition of genes associated with gastric differentiation, epithelial-to-mesenchymal transition (EMT), and metastasis (Clu, Hnf4a, Gkn1, Hmga2, Cldn2). Row normalized expression counts are shown in heatmap. The horizontal lines in each violin in (**f**) represent the 75th percentile, median, and 25th percentile. p values of two-sided Wilcoxon rank-sum test are displayed. **g,** Gene set enrichment analysis using the MSigDB Hallmarks (H1) shows the top 3 enriched gene sets for upregulated (red, log2 fold change > 2 and FDR < 0.05) and downregulated (blue, log2 fold change < −1 and FDR < 0.05) genes in LUAD compared to AdenomaG. Log10 (1/p value) is plotted (one-sided hypergeometric test). EMT = epithelial-to-mesenchymal transition. **h,** Schematic illustration of the snRNA-seq pipeline in *K-MADM-Trp53* lung tumors. The schematic is created with BioRender.com. **i-j,** UMAP plot of single-cell gene expression profiles in *K-MADM-Trp53* lung tumors (n = 6,300 cells). Different cell type clusters identified by unsupervised clustering were labeled with different colors (**i**) based on gene set enrichment patterns of adenoma (n = 1,631) and LUAD (n = 4,669)-specific genes (**j**).

**Extended Data Fig. 7.**
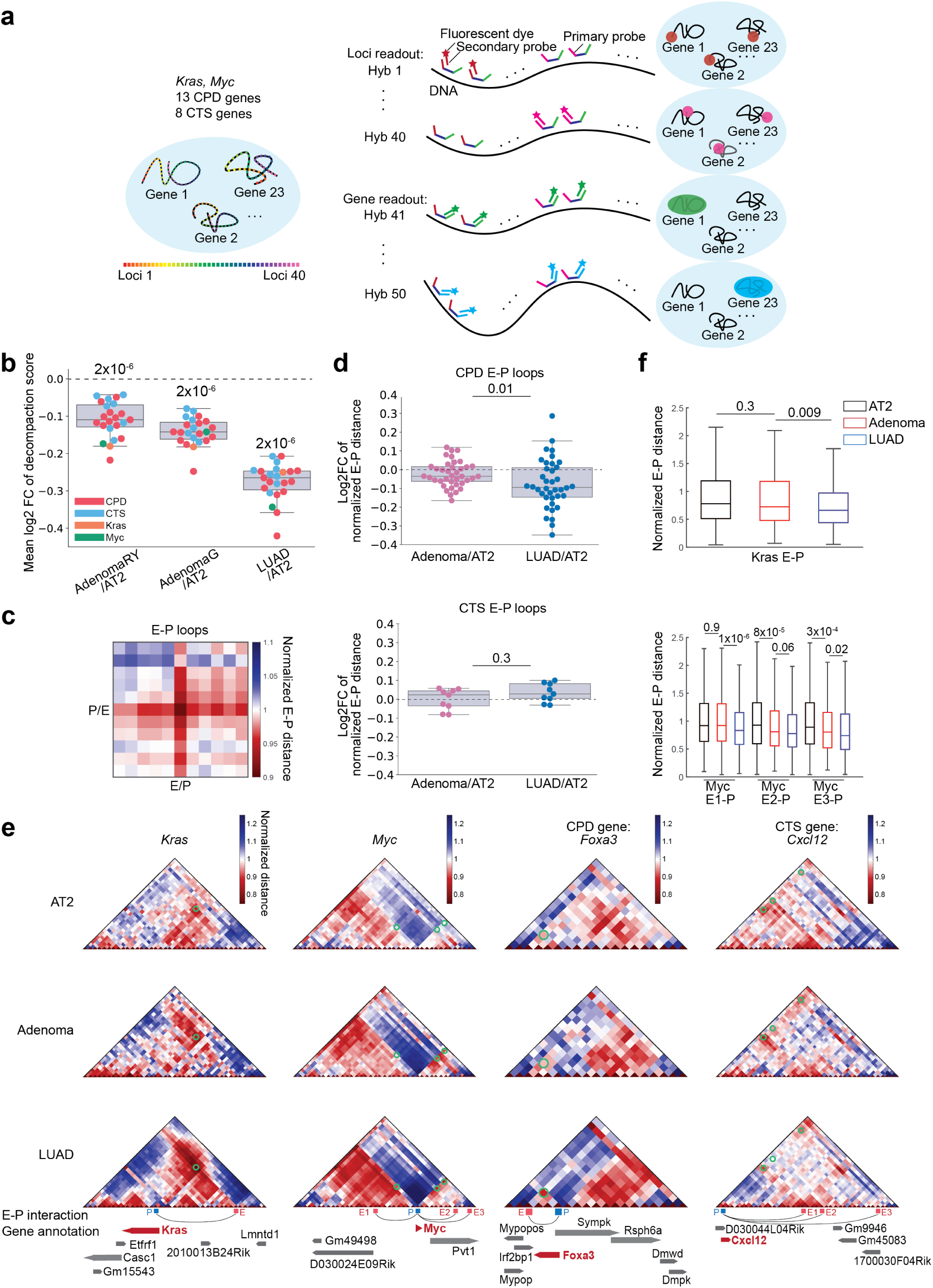
Fine-scale tracing of candidate drivers and suppressors of cancer progression. **a,** Schematic illustration of high-resolution chromatin tracing targeting the cis regulatory regions of 23 genes, including CPD, CTS, *Kras*, and *Myc*. **b,** Distribution of the mean log2 fold change of decompaction (mean inter-loci distance) scores of each target gene, comparing each cancer state to AT2 state. n = 960, 460, 368, 5723 for normal AT2, Adenoma RY, Adenoma G, and LUAD cells. Cell numbers in each cell state are identical in **b-f**. **c,** Pileup heatmap of normalized E-P distances centered around each E-P loop. E-P loops of CPD genes are called in LUAD cells. E-P loops of CTS genes are called in AT2 plus adenoma cells. **d,** Distribution of the log2 fold change of the normalized distances between promoter and putative enhancers of CPD and CTS genes, comparing each cancer state to AT2 cells. **e,** Normalized inter-loci distance matrices of the target genomic regions surrounding the *Kras*, *Myc*, *Foxa3* (CPD), and *Cxcl12* (CTS) genes in AT2, adenoma, and LUAD cell states. The green circles designate putative enhancer-promoter contacts. Putative enhancer and gene annotation tracks are aligned to the target regions. **f,** Distribution of the normalized distances between promoter and putative enhancers of the *Kras* and *Myc* genes.

**Extended Data Fig. 8.**
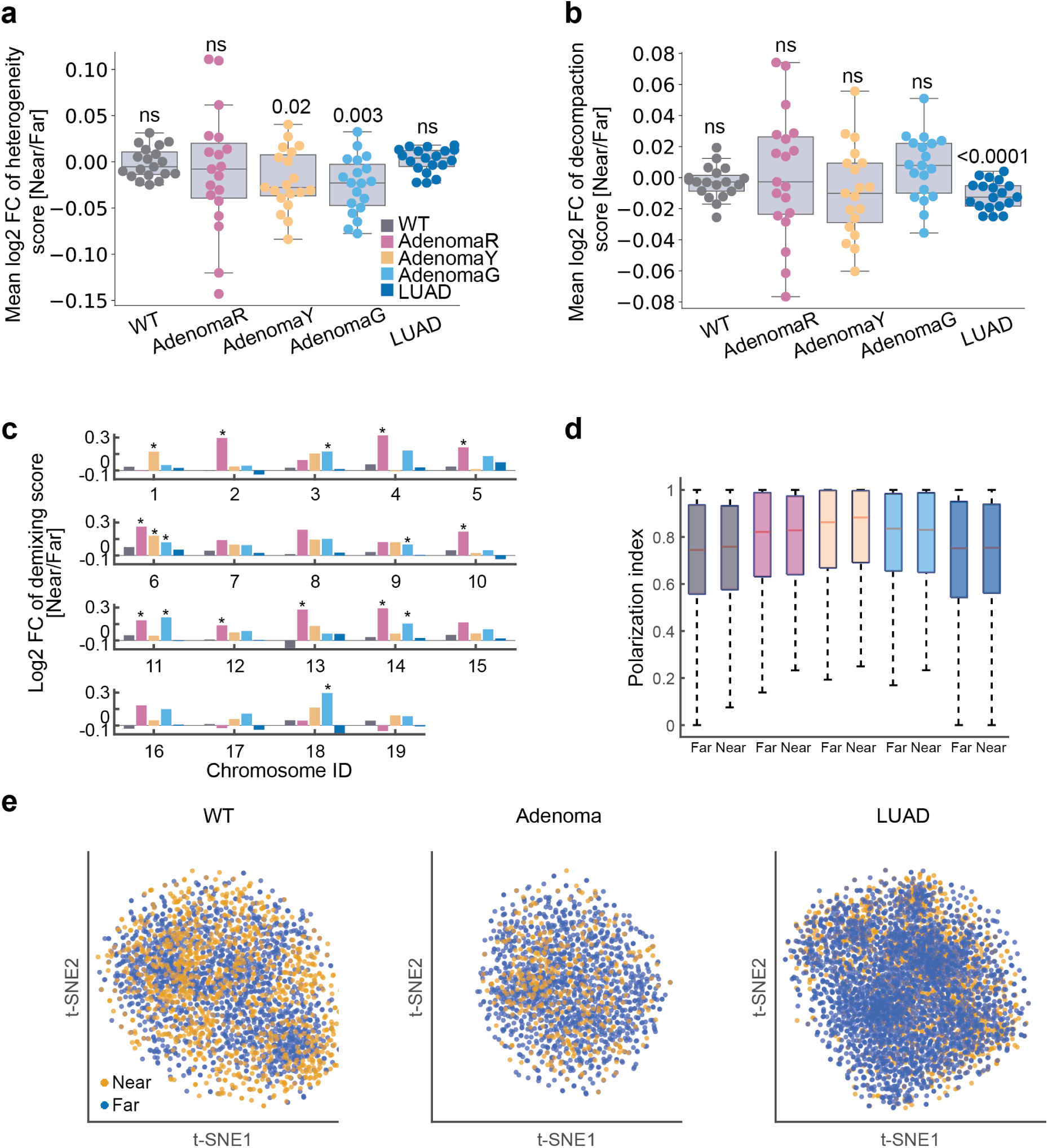
3D genome organization in cancer cells is largely independent of spatial proximity to immune cells. **a-c,** Mean log2 fold change of heterogeneity (**a**), decompaction (**b**), and demixing (**c**) scores of each chromosome, comparing AT2/cancer cells near (less than 10 μm) versus far (more than 10 μm) from CD45+ immune cells. The p values (**a, b**) or FDR (**c**) with significance (< 0.05) from two-sided Wilcoxon signed-rank test (**a, b**) or two-sided Levene’s test (**c**) are displayed. **d,** Distribution of polarization indices of A-B compartments in AT2/cancer cells near or far from immune cells. Two-sided Wilcoxon rank-sum test yielded no significant p values (p < 0.05). **e,** t-SNE plots of single-cell 3D chromatin conformations in AT2, adenoma, and LUAD cells show no distinct clusters based on spatial proximity to immune cells. In (**a, b, d**), the horizontal lines of each box from top to bottom represent the 75th percentile, median, and 25th percentile. Whiskers extend to the non-outlier maximum and non-outlier minimum. Outliers are defined as values at least 1.5 times interquartile range away from the top or bottom of the box.

**Extended Data Fig. 9.**
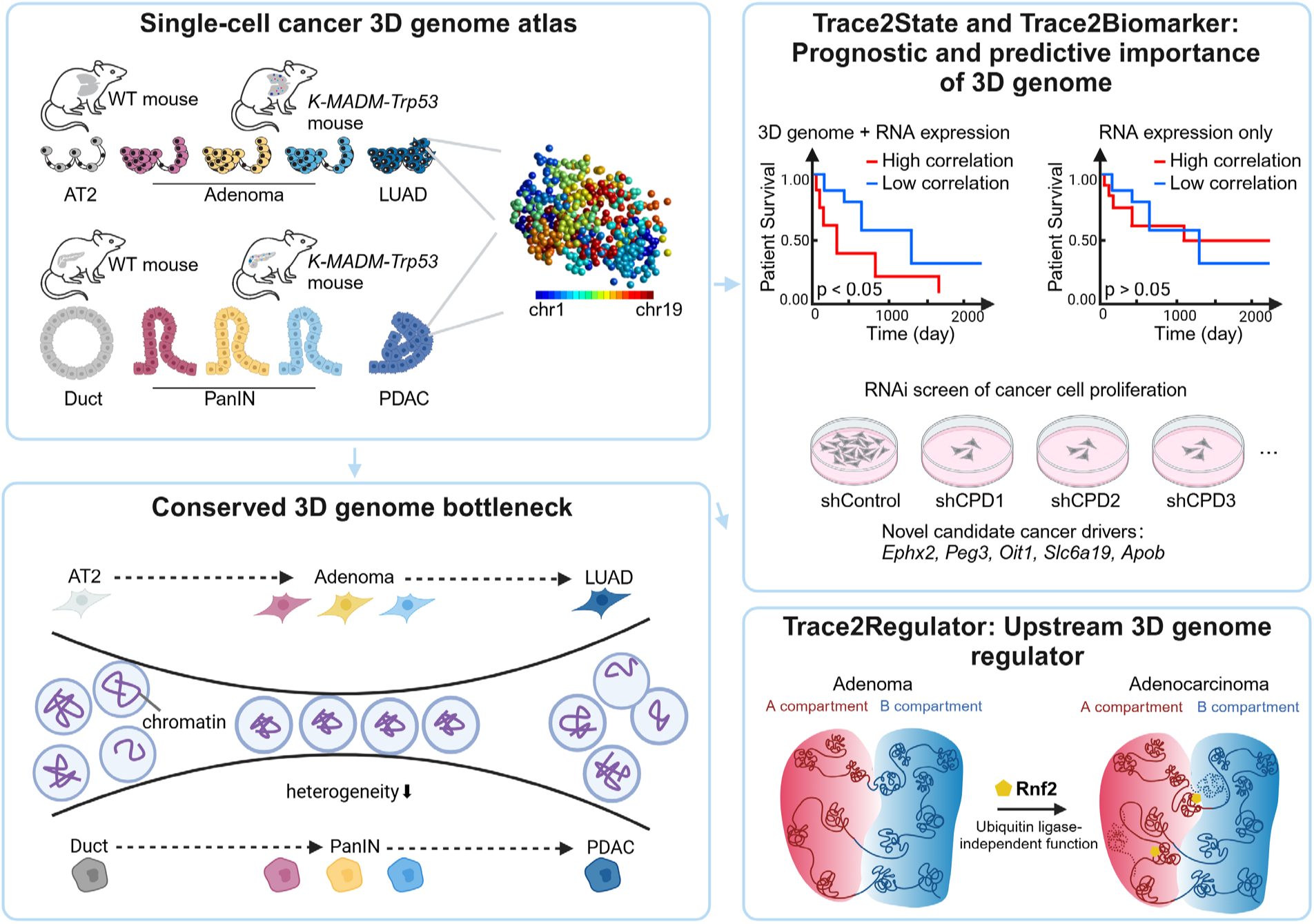
Schematic illustration of the experimental approach and major findings. In this work, we generated single-cell 3D genome atlases during lung and pancreatic cancer progression. Our data revealed stereotypical, stage-specific and conserved alterations in 3D genome folding as cancers progress from normal to preinvasive to invasive tumors, elucidating a potential structural bottleneck during early cancer progression. We developed “Trace2State” and “Trace2Biomarker” pipelines and revealed the utility of 3D genome mapping in discovering prognostic and predictive biomarkers. We further developed a “Trace2Regulator” pipeline and identified a ubiquitin ligase-independent role for Rnf2 in 3D genome regulation.

## Notes

### Summary of Updates

Figures2-6 updated; Supplemental files updated; Title, abstract and author list updated.

